# Microbial Dynamics Across Commercial Spaceflights of Varying Duration

**DOI:** 10.64898/2026.05.29.728784

**Authors:** Aparna Krishnavajhala, Marie-Claude Gingras, Tasha M. Santiago-Rodriguez, Yi Chen, Dilrukshi Bandaranaike, Sravya Bhamidipati, Xiang Qin, Kavya Kottapalli, Zeineen Momin, Abirami Santhanam, Kimberly Walker, Qiaoyan Wang, S. Michelle Griffin, Michal M. Masternak, Matthew C. Ross, Donna Muzny, Jimmy Wu, Emmanuel Urquieta, Richard A. Gibbs, Harsha Doddapaneni

## Abstract

Spaceflight introduces environmental stressors that can alter human microbiomes and immune responses. We analyzed 259 and 17 biospecimens for microbiome and single-cell studies from n=6 astronauts across two commercial ISS missions: Axiom 2 (10-day mission) and Axiom 3 (21-day mission). Samples included saliva, stool, urine, and body swabs from 10 anatomical sites, profiled via 16S ribosomal RNA (rRNA) gene amplicon sequencing. Gut and oral microbiomes remained stable, while skin-associated communities exhibited transient diversity shifts post-flight. Taxonomic analysis revealed individual and site-specific patterns, as well as from the spaceflight/ISS environment. Cytokine profiling from single-cell data indicated immune activation, with IL-32 and IL-16 transcripts elevated in Axiom 2 and Axiom 3. Microbiome and interleukin correlation analyses found 14 significant associations in Axiom 2 data. These findings provide an integrated view of microbiome individuality. Our study represents a descriptive comparison of two missions, informing strategies for crew health on future long-duration missions.

## INTRODUCTION

Spaceflight presents a unique combination of environmental stressors that can significantly impact human physiology, including the microbiome. The current study involved multiple time point sampling of the human microbiome across two spaceflight missions, capturing microbial changes throughout the journey from Earth to the International Space Station (ISS) and back. In space, the crew is exposed to microgravity, cosmic radiation, confinement in enclosed habitats, and sterile or semi-sterile surroundings ^1,2^. These conditions can disrupt the balance and composition of the human microbiome, leading to changes in microbial diversity, abundance, and functionality ^3^. Therefore, understanding how the microbiome responds to these stresses will be important to plan countermeasures that will contribute to maintaining its stability. This is critical for astronaut health and performance, especially on long-duration missions to the Moon or Mars, where medical support and resupply are limited ^4,5^.

The microbiome is essential to human health, contributing to key physiological functions such as nutrient digestion, metabolic processing, immune system modulation, and protection against pathogenic organisms ^6,7^. Interleukins such as IL-32 and IL-16 are among the key immune mediators that regulate inflammation and maintain epithelial barrier integrity, making them critical targets for understanding immune modulation during spaceflight ^8,9^. The microbiome is not uniform across the body but instead exhibits distinct compositions and functions depending on the site. For instance, the gut microbiome is densely populated and primarily involved in metabolism and immune development, the oral cavity supports diverse microbial biofilms relevant to dental and systemic health, the skin microbiome varies by moisture and oil content and contributes to barrier defense, and the urogenital and urinary tracts harbor lower-density but functionally significant communities ^10–12^. This body-site specificity reflects the adaptation of microbial communities to unique environmental niches and is crucial for maintaining localized and systemic homeostasis ^13,14^.

Previous studies have started to uncover the effects of spaceflight on the human microbiome, with some of the notable insights emerging from the NASA Twins Study and other investigations ^4,15,16^. The NASA Twins Study found that the twin who spent nearly a year on the ISS showed changes in microbial diversity and composition compared to his Earth-bound sibling ^15^. Other studies by Voorthies *et al* have shown that long-duration missions can lead to microbial dysbiosis or resilience depending on body site ^4^. Despite these findings, many studies remain constrained by small sample sizes (often one or two individuals ^15^), and are limited to fewer body sites ^4^, or collection time points. Similarly, the Inspiration4 mission conducted a microbiome study using samples from 10 body sites across four individuals during a 3-day orbital flight and reported short-term spaceflight induced transient shifts in skin-associated microbiomes, while gut and oral communities remained largely stable, alongside coordinated changes in immune signaling pathways ^16^. Recent studies have also characterized microbial communities associated with spaceflight habitats, including the International Space Station and China Space Station, highlighting environmental microbial composition, functional potential, and spacecraft-associated ecological dynamics ^17,18^.

While these studies have provided valuable insights into both host-associated and environmental microbiomes in spaceflight contexts, they remain limited in capturing longitudinal data, restricted body-site sampling, and lack integrated immune profiling. Our study characterized the microbiome from two commercial missions with multi-national crew to the ISS, namely the Axiom 2 (Ax-2) mission with a duration of 10 days and the Axiom 3 (Ax-3) mission with a duration of 21 days. Importantly, this study introduces several methodological and conceptual advances relative to prior spaceflight microbiome investigations. Specifically, we leverage multi-site, longitudinal sampling across two independent commercial ISS missions of differing durations (10 and 21 days), enabling a broader characterization of microbiome dynamics across ten anatomical sites in addition to saliva, stool, and urine. Furthermore, the comparative design across missions provides a unique opportunity to examine microbiome and single-cell immune responses during the first three weeks of spaceflight, a relatively understudied period in human space biology. By integrating 16S rRNA gene sequencing with single-cell transcriptomic profiling, this work also offers a combined view of microbial dynamics and host immune activation. Collectively, these features position this study as a hypothesis-generating framework for understanding host–microbiome interactions in confined spaceflight environments. While these missions differ in duration, the study was designed to provide a descriptive comparison across two distinct flight contexts, reflecting combined influences of crew composition, individual variability, and mission environment. Ax-2 involved four astronauts while Ax-3 included two astronauts. This study design enabled the comparison of the microbiome across 10 body sites, saliva, stool and urine samples in n = 6 astronauts, from missions of 10- and 21-day durations, thereby, providing the first preliminary view of shifts in the human microbiome during the first three weeks in spaceflight. This approach also allows us to evaluate subject-specific and site-specific microbial patterns, including differences at the taxonomic level that may contribute to variability in host responses.

Our paper focuses on four main aims: (1) examining microbial dynamics pre and post-flight, (2) investigating bacterial introduction from the Spaceflight/ISS environment, (3) identifying bacterial exclusivity unique to individual participants, and (4) study association between interleukin expression and microbial dynamics. Together, these aims provide a comprehensive understanding of the impact of space travel on microbial dynamics. A companion paper discusses molecular biology and multi-omics results from these two missions ^19^. In addition, this integrative and multi-mission design enables the identification of individualized microbial signatures and their potential association with immune responses, further supporting the hypothesis-generating nature of this study.

Our findings help establish a standardized framework for tracking temporal shifts in microbial relative abundance using 16S ribosomal RNA gene (rRNA) amplicon sequencing across multiple spaceflight time points. Whole-genome sequencing (WGS) was also performed on a sub-set of the biospecimens to validate findings from 16S rRNA gene amplicon sequencing. Single-cell transcriptomics provided complementary insights into interleukin dynamics during spaceflight. Together, these integrative approaches enhance our understanding of immune modulation and microbial resilience in response to spaceflight stressors of varying durations. These findings presented herein are intended to be descriptive and hypothesis-generating, providing a foundation for future larger-scale investigations.

## RESULTS

In this study, samples were collected from two missions, namely Ax-2 and Ax-3, involving a total of n = 6 subjects; n = 4 from Ax-2 and n = 2 from Ax-3. The microbiome study involved 259 samples, while the single-cell study included 17 samples, totaling 276 biospecimens, that were collected from these six individuals.

The collections used for microbiome study comprised four different sample types, including saliva, stool, urine, and body swabs obtained from 10 different locations on the body [Volar Forearm (ARM), Post-Auricular (EAR), Gluteal Crease (GLU), Occiput (NAP), Nasal (NAC), Oral (ORC), Axillary Vault (PIT), Glabella (TZO), Umbilicus (UMB), and Toe Web Space (WEB)], across four time points from each mission. For Ax-2, samples were collected at three pre-launch time points (L-90, L-30 and L-3) and one post-return time point (R+6), and samples collected at L-30, L-3, and R + 6 time points were utilized for 16S rRNA gene amplicon sequencing. For Ax-3, samples collected at two pre-launch time points (L-30 and L-2) and two post-return time points (R+1 and R+21) were utilized in this study. From n = 6 (Ax-2, n = 4 and Ax-3, n = 2), a total of 155 and 104 samples collected from Ax-2 and Ax-3, respectively, resulting in a total of 259 biospecimens) (Figure 1) were included in this study for 16S rRNA gene amplicon sequencing.

**Figure 1.**
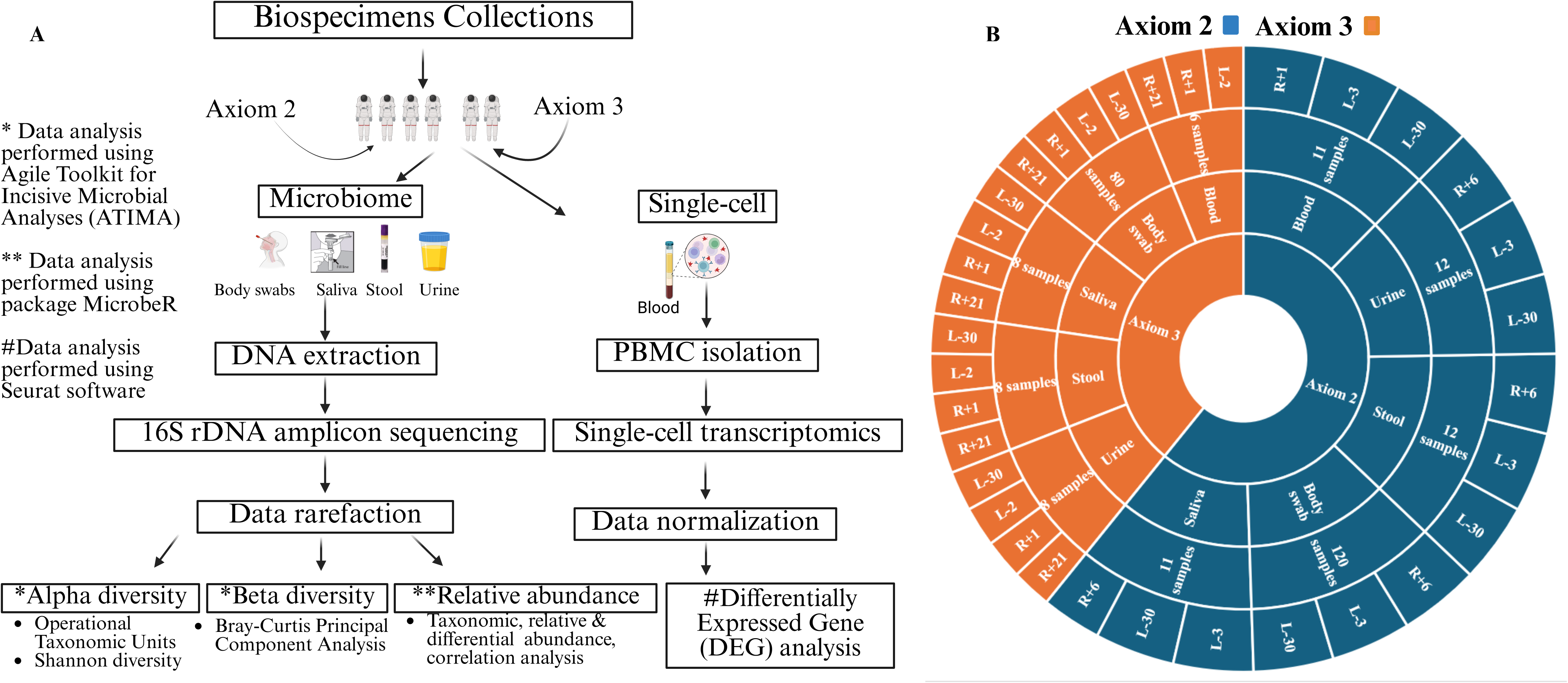
Biospecimen collections and sequencing workflows across two space missions: (A) Schematic representation of the biospecimen collections from Ax-2 (10 day) and Ax-3 (21 day) missions with four and two participants respectively, including stay on the ISS. Four non-blood sample types, body swabs, saliva, stool, and urine, were collected at multiple time points, including pre-flight (2 time points each for Ax-2 and Ax-3) and post-flight (One time point for Ax-2, two for Ax-3). Blood sample was collected at multiple time points, including pre-flight (Two time points for Ax-2 and one for Ax-3) and post-flight (one time point for Ax-2, and two for Ax-3). DNA was extracted from the 259 biospecimens collected from the 6 individuals; the V4 region of the 16S rRNA gene was sequenced. Data rarefied based on sample type and microbial diversity, and abundance, were evaluated. PBMCs were isolated from 18 blood-derived biospecimens, of which 17 were successfully processed for single-cell RNA sequencing and downstream transcriptomic analyses. (B) Circular diagram showing sample distribution by mission (Axiom-2 in blue, Axiom- 3 in orange), sample type and their numbers, and collection time points (L = pre-flight, R = post-flight) from which 16S rRNA gene amplicon and scRNA sequence data were generated.

This resulted in a multi-time point and overlapping sampling design. The crew spent 10 and 21 days on the ISS for Ax-2 and Ax-3 missions, respectively (Figure 1). This longitudinal and multi-site sampling strategy represents a unique study design for evaluating early microbiome and its association with interleukins compared to post-flight in human spaceflight.

### Read counts per sample from 16S rRNA gene amplicon sequencing

For the Ax-2 cohort, of the 155 biospecimens processed, one sample failed during the initial quality control step and was excluded from sequencing. Additionally, 12 samples (one saliva and 11 body swab samples) produced insufficient mapped reads and failed QC. Sequencing data was successfully generated for the remaining 142 biospecimens, corresponding to a success rate of 91.61%. For the Ax-3 cohort, of the 104 biospecimens processed, one (urine) sample produced insufficient mapped reads and failed QC. Sequencing data was successfully generated for the remaining 103 biospecimens, corresponding to a success rate of 99%. 16S rRNA gene amplicon sequencing was favored over whole-genome sequencing (WGS) because it is lower in cost, requires less DNA input, and is supported by standardized analysis pipelines. Data also results in taxonomic profiling similar to WGS and making it easier to conduct cross-study comparisons involving both low and high biomass samples.

The average, minimum, and maximum number of sequencing reads, and their rarefaction cut-off for each sample type for both Ax-2 and Ax-3 missions are summarized in Table 1, which also shows the rarefaction values used, and the standard deviation (SD). Given the diverse nature and biomass of the collected samples, differing rarefaction values were utilized prior to analysis to minimize disparate results due to differing sequencing depths. Saliva, stool, urine, and body swab samples were rarefied to 61,820; 305; 5,930 and 192 reads for the Ax-2 mission, and 22,056; 33,799; 1,134 and 6,131 reads for the Ax-3 mission, respectively (Table 1). Body swab samples were also rarefied on a per-sample type basis to minimize disparate results due to the differences in sequencing depth for both the Ax-2 and Ax-3 mission datasets (Table 1). One saliva and 11 body swab samples from the Ax-2 mission, and one urine sample from the Ax-3 mission were dropped from the analysis due to an insufficient number of reads.

**Table 1.**
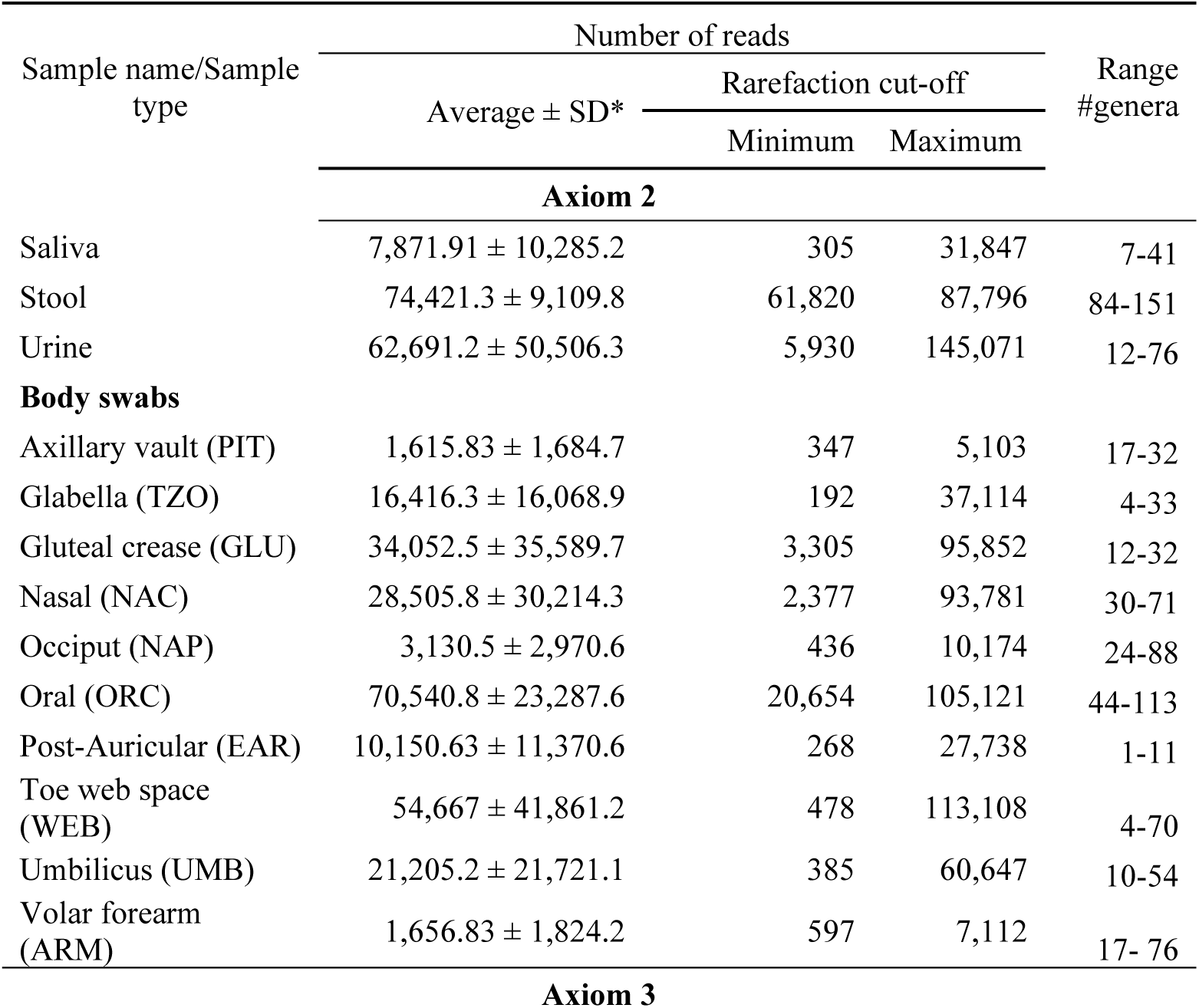
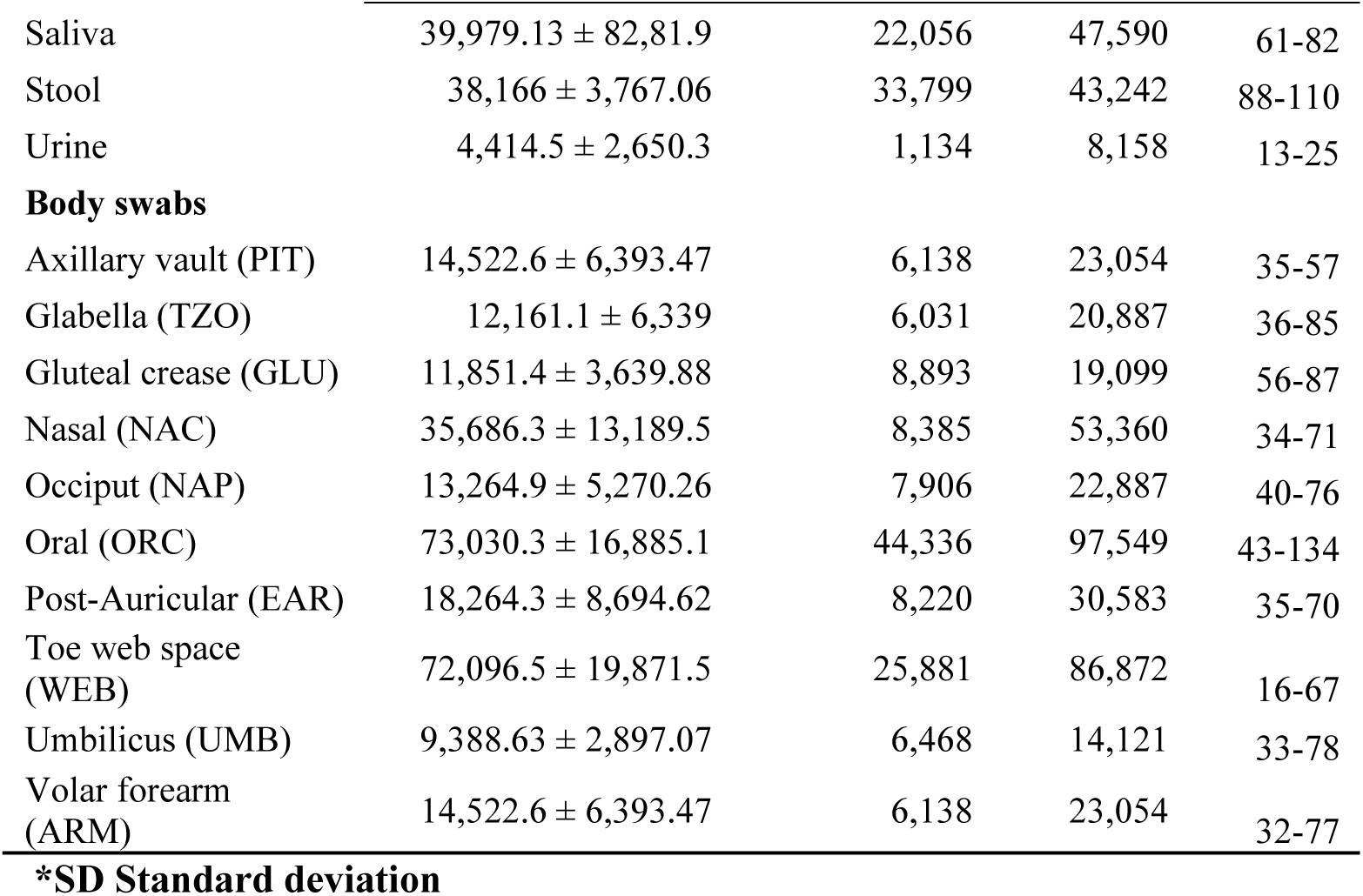
Sample statistics. Table shows the average, minimum, and maximum number of sequencing reads, standard deviation, and rarefaction cut-off for saliva, stool, urine, and body swab samples after 16S rRNA gene amplicon sequencing. The minimum and maximum number of microbial genera from each sample type for Ax-2 and Ax-3 missions are also shown. Rarefaction was performed on a per-sample type basis to minimize the effect of differing rarefaction cutoff values on diversity and taxonomic profiles.

| Sample name/Sample type | Number of reads |  |  | Range #genera |
| --- | --- | --- | --- | --- |
| | Average ± SD* | Rarefaction cut-off | | |
|  |  | Minimum | Maximum |  |
| Axiom 2 |  |  |  |  |
| Saliva | 7,871.91 ± 10,285.2 | 305 | 31,847 | 7-41 |
| Stool | 74,421.3 ± 9,109.8 | 61,820 | 87,796 | 84-151 |
| Urine | 62,691.2 ± 50,506.3 | 5,930 | 145,071 | 12-76 |
| Body swabs |  |  |  |  |
| Axillary vault (PIT) | 1,615.83 ± 1,684.7 | 347 | 5,103 | 17-32 |
| Glabella (TZO) | 16,416.3 ± 16,068.9 | 192 | 37,114 | 4-33 |
| Gluteal crease (GLU) | 34,052.5 ± 35,589.7 | 3,305 | 95,852 | 12-32 |
| Nasal (NAC) | 28,505.8 ± 30,214.3 | 2,377 | 93,781 | 30-71 |
| Occiput (NAP) | 3,130.5 ± 2,970.6 | 436 | 10,174 | 24-88 |
| Oral (ORC) | 70,540.8 ± 23,287.6 | 20,654 | 105,121 | 44-113 |
| Post-Auricular (EAR) | 10,150.63 ± 11,370.6 | 268 | 27,738 | 1-11 |
| Toe web space (WEB) | 54,667 ± 41,861.2 | 478 | 113,108 | 4-70 |
| Umbilicus (UMB) | 21,205.2 ± 21,721.1 | 385 | 60,647 | 10-54 |
| Volar forearm (ARM) | 1,656.83 ± 1,824.2 | 597 | 7,112 | 17- 76 |
| Axiom 3 |  |  |  |  |
| Saliva | 39,979.13 ± 82,81.9 | 22,056 | 47,590 | 61-82 |
| Stool | 38,166 ± 3,767.06 | 33,799 | 43,242 | 88-110 |
| Urine | 4,414.5 ± 2,650.3 | 1,134 | 8,158 | 13-25 |
| **Body swabs** |  |  |  |  |
| Axillary vault (PIT) | 14,522.6 ± 6,393.47 | 6,138 | 23,054 | 35-57 |
| Glabella (TZO) | 12,161.1 ± 6,339 | 6,031 | 20,887 | 36-85 |
| Gluteal crease (GLU) | 11,851.4 ± 3,639.88 | 8,893 | 19,099 | 56-87 |
| Nasal (NAC) | 35,686.3 ± 13,189.5 | 8,385 | 53,360 | 34-71 |
| Occiput (NAP) | 13,264.9 ± 5,270.26 | 7,906 | 22,887 | 40-76 |
| Oral (ORC) | 73,030.3 ± 16,885.1 | 44,336 | 97,549 | 43-134 |
| Post-Auricular (EAR) | 18,264.3 ± 8,694.62 | 8,220 | 30,583 | 35-70 |
| Toe web space (WEB) | 72,096.5 ± 19,871.5 | 25,881 | 86,872 | 16-67 |
| Umbilicus (UMB) | 9,388.63 ± 2,897.07 | 6,468 | 14,121 | 33-78 |
| Volar forearm (ARM) | 14,522.6 ± 6,393.47 | 6,138 | 23,054 | 32-77 |

**\*SD Standard deviation**

### Alpha diversity analysis

To assess alpha diversity or within-sample microbial diversity, Observed Operational Taxonomic Units (OTUs) and the Shannon Diversity Index of stool, saliva, urine, and body swab samples from six subjects (four from Ax-2 and two from Ax-3) were calculated (Figures 2A–B). Observed OTUs is a measurement of species richness, while the Shannon index incorporates both richness and evenness. Significant differences in alpha diversity were assessed through non-parametric Mann-Whitney U test or Kruskal-Wallis for the comparison of 2 groups or 3 or more groups, respectively, and p-values were adjusted using FDR correction. Boxplots were generated for each mission and colored by time points: L-30, L-3, and R+6 for Ax-2; L-30, L-2, R+1, and R+21 for Ax-3 (Figures 2A–B; Figures S1–2). These visualizations highlight temporal trends in richness and evenness across body sites.

**Figure 2.**
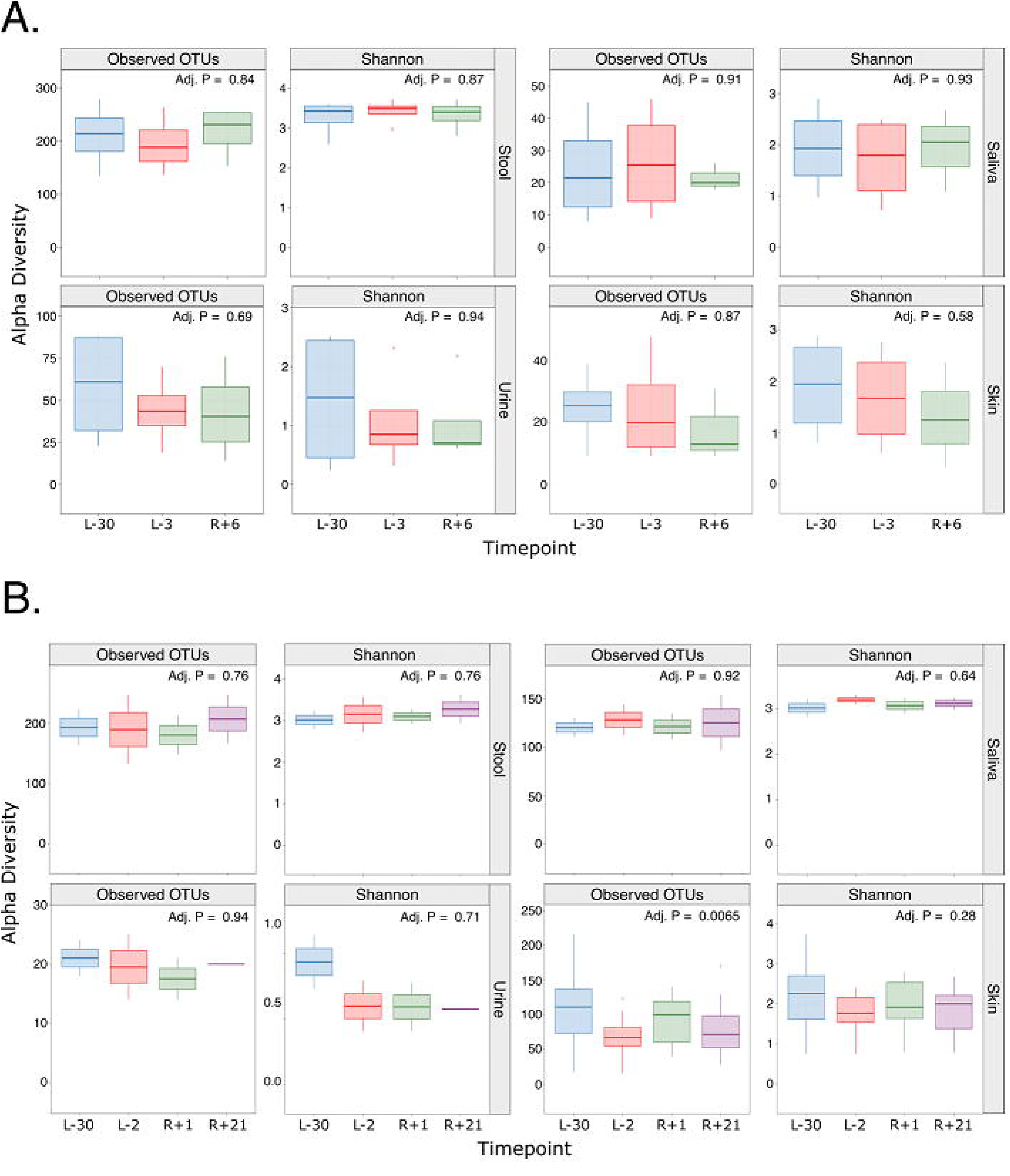
Alpha diversity analysis of microbiome (stool, saliva, urine and body swab samples) from Ax-2 (A) and 3 (B) participants. Panel A and B show the number of observed Operational Taxonomic Units (OTUs) and the Shannon diversity index for stool, saliva, urine, and body swab samples that were rarefied based on sample type for the Ax-2 and Ax-3 missions, respectively. Boxplots depict changes in the number of OTUs and Shannon diversity of the human microbiome. Box plots are colored by time points, namely L-30, L-3 and R+6 for the Ax-2 mission, and L-30, L-2, R+1, and R+21 for the Ax-3 mission. For Ax-2, a total of 12 stool, 11 saliva, 12 urine and 120 body swab samples were included across all time points. For Ax-3, a total of 8 stool, 8 saliva, 8 urine and 80 body swabs samples were included across all time points. Significant differences were assessed through non-parametric Mann-Whitney U test or Kruskal-Wallis for the comparison of 2 groups or 3 or more groups, respectively, and p-values were adjusted using FDR correction. Adjusted p-values are shown on the plots.

Stool alpha diversity remained highly stable across both Ax-2 (observed OTUs adj. P = 0.84; Shannon diversity adj. P = 0.87) and Ax-3 (adj. P = 0.76 for both metrics) missions. Saliva exhibited slight, non-significant decreases in alpha diversity post-flight in both Ax-2 (observed OTUs adj. P = 0.91; Shannon adj. P = 0.93) and Ax-3 (observed OTUs adj. P = 0.92; Shannon adj. P = 0.64) missions. Urine displayed low richness compared to stool, with minimal variation in both Ax-2 (observed OTUs adj. P = 0.69; Shannon = 0.94) and Ax-3 (observed OTUs adj. P = 0.94; Shannon = 0.71) (Figure 2A and 2B).

Body swabs exhibited the largest changes in alpha diversity in association with spaceflight (Figure S1). For instance, a significant increase in alpha diversity was observed in occiput (NAP) samples collected at time point R+6 during the Ax-2 mission (observed OTUs adj. P = 0.024; Shannon adj. P = 0.05). Significant shifts in Shannon diversity were observed in oral (ORC) samples during the Ax-2 mission (adj. P = 0.024) (Figure S1). In Ax-3, a significant increase in richness post-flight (observed OTUs adj. P = 0.0065) was noted without corresponding changes in evenness (Shannon adj. P = 0.28). Site-specific trends included Ax-2 sensitivity at NAP and ORC, and Ax-3 trends at glabella (TZO) (observed OTUs adj. P = 0.11), axillary vault (PIT) (Shannon adj. P= 0.13), and gluteal crease (GLU) (observed OTUs adj. P = 0.14). No significant differences were noted in alpha diversity for body swab samples collected during the Ax-3 mission (Figure S2).

Overall, alpha diversity analysis showed that stool and saliva microbiomes remained stable across both missions over time (Figure 2A and 2B). Notably, urine samples exhibited consistently low diversity compared to stool samples, reflecting their low-biomass nature. In contrast, body swab samples demonstrated the greatest sensitivity to spaceflight, with Ax-2 showing significant richness and evenness changes at sites including NAP and ORC (Figure 2A), while Ax-3 exhibited increased species richness post-flight (observed OTUs adj P = 0.0065), indicating higher richness immediately after return compared to pre-flight (Figure 2B). Shannon diversity did not exhibit significant changes (adj. P = 0.28), demonstrating that, while richness increased, community evenness remained relatively stable (Figure 2B).

### Beta diversity analysis

Beta diversity of six subjects (four from Ax-2 and two from Ax-3) was assessed through Bray-Curtis distances and visualized through Principal Coordinate Analyses (PCoA) plots. Significance was assessed through a Permutational Multivariate Analysis of Variance (PERMANOVA). Results revealed distinct microbial community differences between samples, as shown in the PCoA plots for both Ax-2 (Figure 3A) and Ax-3 (Figure 3B) missions. These patterns were based on Permutational Multivariate Analysis of Variance (PERMANOVA) testing.

**Figure 3.**
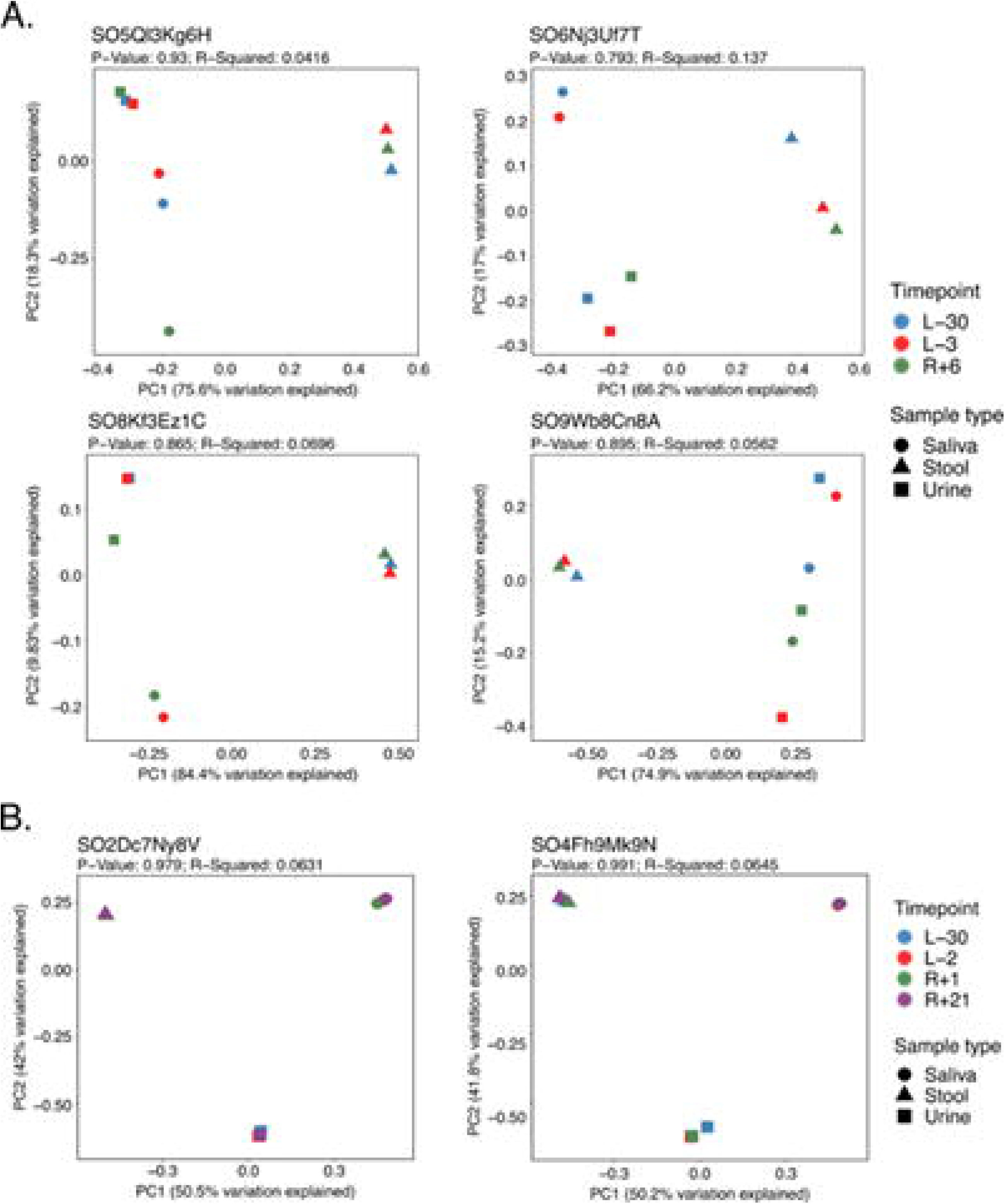
Beta diversity analysis. Beta diversity analyses were assessed through Bray-Curtis distances and visualized through Principal Coordinate Analyses (PCoA) plots. Significance was assessed through a Permutational Multivariate Analysis of Variance (PERMANOVA). PCoA plots of the Bray-Curtis distances of stool, saliva, and urine samples across subjects and time points in Ax-2 (A) and Ax-3 (B) missions are shown here. P-values were calculated using PERMANOVA and are displayed on the plot with the R-Squared value.

Across both missions and subjects, samples clustered strongly by sample type rather than time point, indicating that body site identity was the dominant driver of microbial composition. No clear temporal separation was observed for saliva, stool, or urine, suggesting stability of these niches under spaceflight conditions. PERMANOVA results confirmed this pattern: Ax-2 subjects showed P-values ranging from 0.793 to 0.93, with R² values ranging from 0.0416 to 0.137, and Ax-3 subjects showed P-values ranging from 0.979 to 0.991, with R² values ranging from 0.0631 to 0.0645, highlighting minimal variance explained by time point.

Within body swabs, NAC (P = 0.015; R² = 0.394) and ARM (P = 0.022; R² = 0.276) samples collected during Ax-2 mission exhibited site-specific post-vs. pre-flight separation, supporting localized shifts in skin-associated microbiota (Figure S3). In contrast, Ax-3 showed no statistically significant changes at any site, though visual clustering by subject was observed, reflecting strong individual signatures rather than temporal effects (Figure S4).

To further evaluate temporal microbial dynamics independent of body site-associated clustering, additional PCoA analyses were generated with biological sample types displayed as separate facets with (Figure S5). This approach enabled visualization of longitudinal microbial variation within saliva, stool, urine, and skin-associated microbiomes individually, thereby reducing the dominant separation driven by anatomical site differences observed in the combined analyses. Faceted ordinations demonstrated that microbial community structure remained largely individualized within each sample category, with only modest temporal shifts observed across mission time points.

### Taxonomic analyses

Relative abundances from six subjects (four from Ax-2 and two from Ax-3) were estimated as the proportion of reads assigned to each taxon within each sample and were used for taxonomic analysis. Taxonomic analysis revealed both subject-specific and mission-dependent patterns across all sample types (Figure 4A–D). For instance, saliva samples from the Ax-2 mission were dominated by *Staphylococcus*, with abundances ranging from 8.52% to 86.23%, followed by *Corynebacterium* with abundances ranging from 2.95% to 69.18%. Subject-specific patterns in relative abundances were observed for specific subjects. For instance, saliva samples from subject SO5Ql3Kg6H shifted from *Staphylococcus* (86.23% at L-30) to *Cellulomonas* (22.95% at R+6). In contrast, saliva samples collected from Ax-3 were dominated by *Streptococcus* (8.12%– 21.77%), *Neisseria* (10.41%–14.34%), *Haemophilus* (2.64%–21.94%), and *Prevotella_7* (5.18%– 19.41%) across all time points.

**Figure 4.**
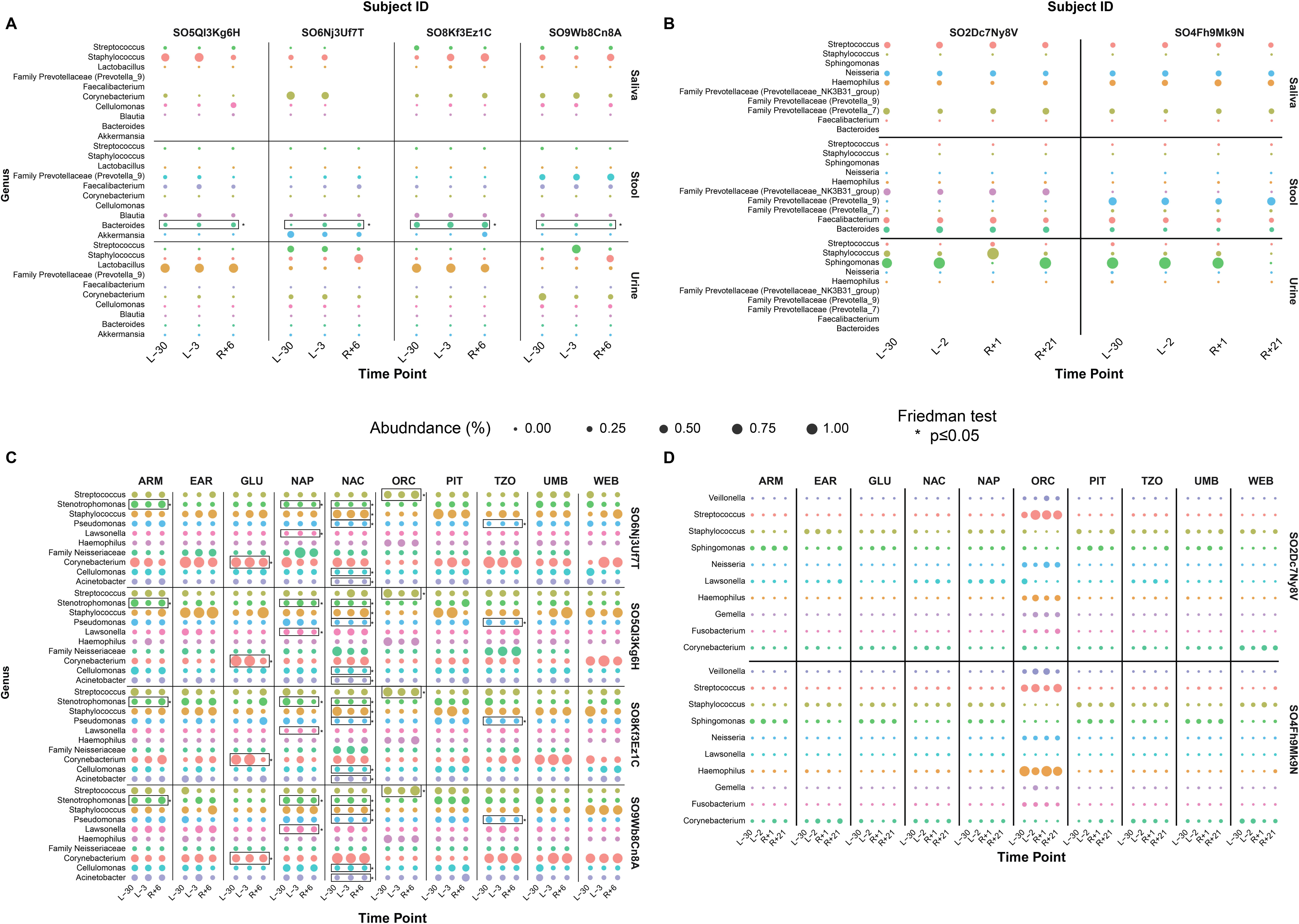
Multi-dimensional analysis of the top 10 most abundant bacterial genera across sample types and missions. Panels A and B show bubble plots for Ax-2 (A) and Ax-3 (B) saliva, stool, and urine samples, illustrating relative abundance of the top 10 genera across biological sample types. Panels C and D depict Ax-2 (C) and Ax-3 (D) body swab samples collected from 10 anatomical sites (ARM, EAR, BLU, NAP, NAC, ORC, PIT, TZO, UMB, WEB). The x-axis represents time points (Ax-2: L-30, L-3, R+6; Ax-3: L-30, L-2, R+1, R+21), and the left y-axis lists bacterial genera. Bubble size corresponds to percentage abundance (0.00–1.00), while the right y-axis distinguishes sample types for A and B panels. These visualizations enable direct comparison of microbial community profiles across time points, sample types, and subjects, highlighting site-specific and mission-dependent patterns in microbiome composition.

Stool microbiomes exhibited subject-specific temporal variation across both missions, while generally remaining dominated by common gut-associated bacterial genera. For instance, stool samples collected from Ax-2 exhibited higher relative abundances of *Bacteroides*, *Prevotellaceae*, and *Faecalibacterium*. Urinary microbiomes also showed potential mission- and subject-specific patterns. For instance, in Ax-2, the urinary microbiomes of two subjects were dominated by *Lactobacillus*, while in the remaining two subjects, *Staphylococcus* and *Corynebacterium* were the predominant bacteria. In contrast, Ax-3 urine samples were primarily dominated by *Sphingomonas*, with intermittent surges of *Staphylococcus* and an occasional presence of *Streptococcus*.

Body swabs across both missions (Figure 4C-D) were enriched for *Corynebacterium* and *Staphylococcus*, with Ax-3 additionally showing *Lawsonella* and *Sphingomonas*. Site-specific trends included Ax-2 TZO samples dominated by *Corynebacterium* and Ax-3 NAP samples enriched for *Lawsonella* ORC samples consistently featured oral taxa, with *Streptococcus* reaching relative abundances of 51% and *Haemophilus* up to 33% at post-flight time points. Opportunistic genera such as *Micrococcus*, *Paracoccus*, and *Comamonas*) appeared exclusively in post-flight samples.

In summary, taxonomic profiling across saliva, stool, urine, and body swab samples revealed strong individuality and sample-type-specific microbial signatures, with dominant genera varying by mission and subject.

To evaluate temporal changes in microbial community composition, genus-level differential abundance analyses from six subjects (four from Ax-2 and two from Ax-3) were performed using MaAsLin3 separately for each mission and body site (Figure S6). Significant time point-associated taxa were identified in selected body sites (q-value < 0.3), including NAP, ORC, stool, and urine samples from Ax-2, and GLU, NAC, TZO, and urine samples from Ax-3. Notably, many of the taxa identified through MaAsLin3 corresponded to dominant genera highlighted in the taxonomic analyses. For example, oral cavity samples exhibited significant associations involving *Streptococcus* and *Haemophilus*.

In addition, complementary non-parametric Friedman tests, followed by paired Wilcoxon post hoc comparisons, further supported these results by identifying limited statistically significant temporal changes across sample types.

### Potential post-flight emergence

We noticed a rare occurrence of potential post-flight emergence of bacteria from the ISS environment. Specifically, in the Ax-3 mission, subject SO4Fh9Mk9N body swabs revealed *Oligella* in ARM (0.52%) at R+1 time point, while it was absent in pre as well as post-flight biospecimens from all other participants. This points to potential post-flight emergence of bacteria, likely from ISS, given their multiple day stay. We followed up this finding with WGS metagenomic sequencing and were able to confirm that in the WGS data, in the preflight biospecimen, there was one read mapped to *Oligella,* while 2,904 reads were identified in the R+1 biospecimen (Table S2). Strain level analysis of WGS data indicated the presence of *Oligella urethralis and O. ureolytica* with bacterial read counts 0 and 1 at L-2; 1,856 and 831 at R+1 respectively, from the same individual (SO4Fh9Mk9N), further supporting the possible post-flight bacterial emergence from the ISS environment.

### Bacterial exclusivity

Our study revealed distinct patterns of potential bacterial taxa that seem to be distinct across individuals and sample types, suggesting personalized microbial signatures to spaceflight. Saliva samples showed *Anaeroglobus* exclusively in subject SO2Dc7Ny8V. Stool samples further underscored the individuality of microbial profiles, where subject SO2Dc7Ny8V contained *Anaeroplasma, Desulfovibrio, and Holdemanella*, while SO4Fh9Mk9N uniquely harbored *Prevotella_7*, *Rikenellaceae_RC9_gut_group*, and *Granulicatella*. Body swabs revealed *Holdemanella* only in SO2Dc7Ny8V, detected in TZO (0.02%) and NAP (0.03%) at L-30, and in additional sites at L-30 (0.25%), R+1 (0.01%), and R+21 (0.17%).

### Interleukin Expression and Microbiome Association in Ax-2

We included 11 blood samples for single-cell data analysis (Figure 1). Analysis of interleukin expression at group level (grouping time points L-30, L-3, R+1 from all 4 individuals) in Ax-2 (Figure 5A) revealed elevated transcript levels of IL-32 and IL-16 at the post-flight time point (R+1), supporting a heightened immune response following spaceflight.

**Figure 5.**
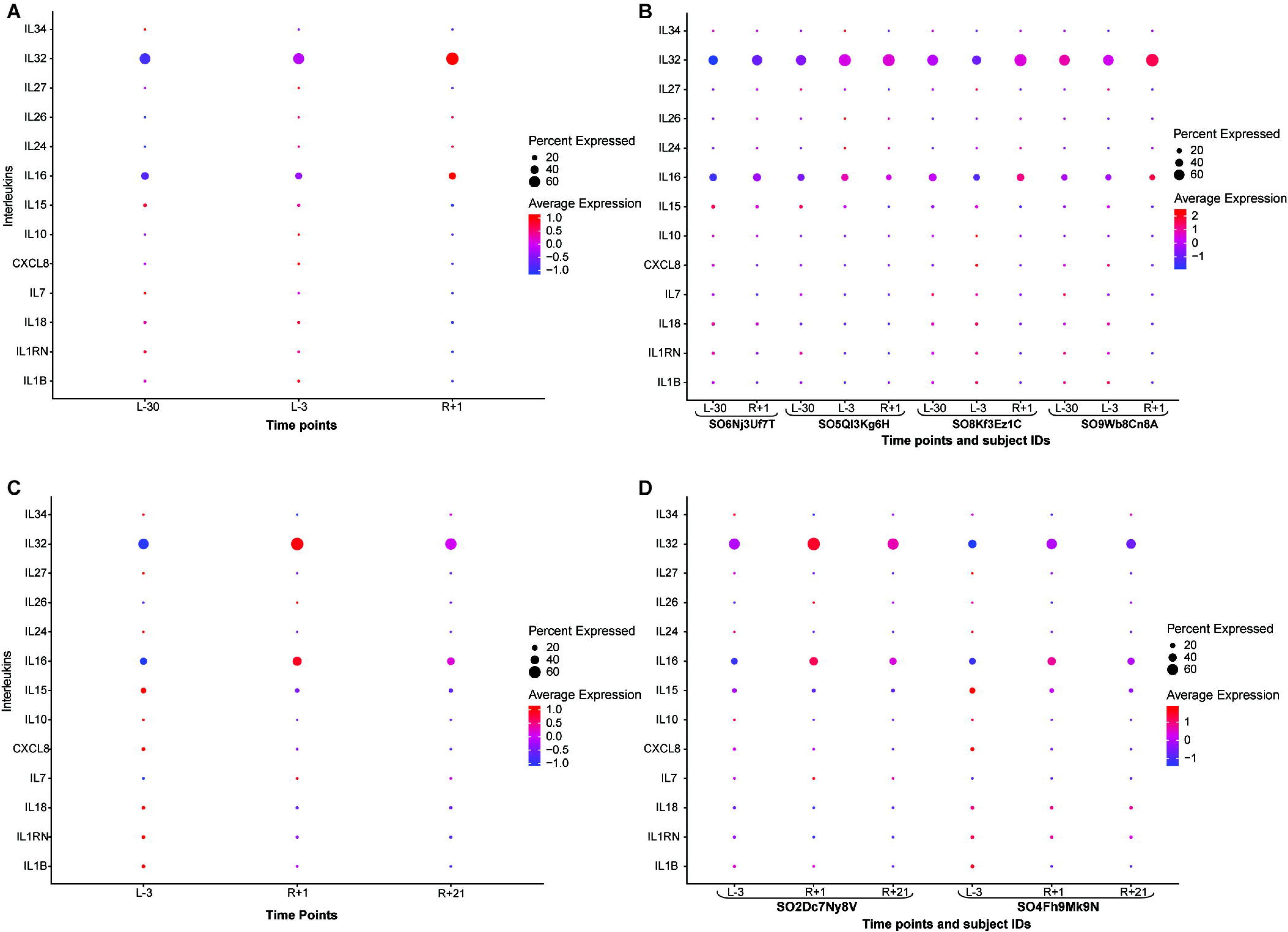
Interleukin expression profiles across missions and time points. Panels A and B show Ax-2 data: (A) group-level interleukin transcript expression across three time points (L-30, L-3, R+1), with colored dots indicating average expression (red = higher [value = up to 2], blue = lower [value = −1]) and dot size representing the percentage of samples expressing each interleukin; (B) subject -level data (n = 4) across the same time points, with subject IDs on the X-axis and interleukins on the Y-axis using the same color and size scheme. Panels C and D show Ax-3 data: (C) group-level transcript expression across three time points (L-2, R+1, R+21) using the same encoding; (D) subject-level data (n = 2) across the same time points, with subject IDs and interleukin subset displayed as in Panel B. These visualizations highlight temporal and subject-specific patterns in interleukin expression.

The coordinated rise in IL-32 and IL-16 may reflect host adaptation to environmental microbial exposure, reinforcing the importance of monitoring immune-microbiome dynamics during space missions. The study also reinforces that subject-level analysis (individual) of interleukin expression from Ax-2 (Figure 5B) across time points L-30, L-3, and R+1 revealed dynamic immune responses associated with spaceflight. Notably, IL-32 and IL-16 showed elevated expression at the post-flight time point (R+1), with IL-32 consistently across all the individuals, while IL-16 showed increased average expression across three individuals (SO6Nj3Uf7T, SO8Kf3Ez1C, and SO9Wb8Cn8A) at post-flight time points. These cytokines are known to mediate inflammatory responses and epithelial barrier regulation.

### Interleukin Expression and Microbiome Association in Ax-3

We included 6 blood samples for single-cell data analysis (Figure 1). Interleukin analysis in Ax-3 (Figure 5C) revealed increased expression in IL-16, and IL-32 and decreased expression of IL-15 at time point R+1 compared to pre-flight time point (L-2), at both group and individual level, supporting immune activation immediately after spaceflight (Figure 5C-D). IL-16 and IL-32 responses declined by R+21, showing a potential return towards baseline activity. Subject-level trends (Figure 5D) further support this pattern.

To better contextualize the contribution of IL-16 and IL-32, we performed microbiome-interleukin correlation analyses of the four subjects integrating Axiom-2 microbial profiles with cytokine transcript expression responses across multiple sample types. A total of 14 significant correlations were identified, involving 13 unique bacterial taxa distributed across five sample types: nasal (NAC), neck (NAP), oral (ORC), stool, and urine. Positive associations (p<0.05) included, ORC (6), stool (3), NAC (1), NAP (1), and urine (1) (Table S3), indicating that oral and, to a lesser extent, gut and nasal niches showed the strongest detectable relationships with cytokine transcript expression levels in this dataset. These positive associations (Pearson r > 0.95, with concordant Spearman results) were observed between specific microbial taxa and IL-16 or IL-32, including *Streptococcus* (NAP) with IL-32 (r = 0.999, p = 0.001), *Corynebacterium* (urine) with IL-32 (r = 0.992, p = 0.008), and multiple oral taxa (Ex: *Cardiobacterium, Rothia, Lautropia*) with IL-16 (Table S3). The subject-level correlations of the above are shown in Figure S7-S9. Overall, these findings suggest that mucosal sites (oral and nasal) and the gut microbiome may represent locations where measurable immune-microbiome associations were observed, with IL-16 and IL-32 showing consistent positive associations across multiple taxa and sample types (Supplementary Table 3). For Axiom 3 (n = 2), correlation analysis was not performed. The temporal alignment between cytokine activation and microbial acquisition supports the hypothesis that spaceflight induces coordinated changes in immune signaling and microbiome composition, underscoring the need to monitor host-microbe interactions during extended missions.

## DISCUSSION

The Ax-2 (10 days) and Ax-3 (21 days) missions provided a unique opportunity to examine microbiome dynamics during short-duration commercial spaceflights of different durations. From analyzing 259 biospecimens from n = 6, this study observed conserved, yet individualized microbial shifts across saliva, stool, urine, and body swabs, suggesting selective pressures from spaceflight. Notably, stool and saliva microbiomes remained highly stable, underscoring the resilience of these niches and aligning with terrestrial evidence of gut and oral community robustness.^4,14^. Notably, this study represents one of the first efforts to integrate multi-site microbiome profiling across multiple commercial ISS missions with concurrent immune transcriptomic data, allowing for a more comprehensive assessment of early host–microbiome responses to spaceflight. In contrast to recent studies focusing on environmental microbiomes within spaceflight habitats ^17,18^, our study specifically examines longitudinal host-associated microbiome dynamics across multiple body sites in astronauts, providing complementary insights into human-specific responses to spaceflight.

Our study is the first to demonstrate microbiome dynamics focused on the first three weeks of spaceflight, expanding on prior works of the year-long NASA Twins Study ^15^ and the 3-day Inspiration4 mission ^16^. While long-duration missions have reported compositional shifts under prolonged exposure ^15^, the short-duration missions like Inspiration4 observed transient skin microbiome changes without major gut or oral shifts ^16^. In another study, Voorhies *et al*. investigated gut microbiome changes during extended ISS stays, focusing on a limited number of body sites, and reported compositional shifts in gut microbiome, and changes in skin microbiome ^4^. Our study examined two commercial ISS missions lasting 10 and 21 days, profiling six astronauts across ten anatomical sites and multiple sample types. Our results revealed sustained variability in skin-associated taxa and delayed recovery in gut communities, even as gut and oral microbiomes remained largely stable, highlighting that skin niches are particularly sensitive to spaceflight conditions.

Indeed, beta diversity analyses demonstrated that individual identity and body site were the primary drivers of microbial community structure across the dataset, whereas spaceflight-associated temporal effects accounted for a comparatively modest proportion of the observed variance. These findings suggest that astronaut microbiomes remained largely individualized and body site-specific throughout the study period, with longitudinal microbial shifts occurring within the context of relatively stable baseline community structures.

Skin-associated microbiomes showed the greatest variability under spaceflight conditions, consistent with prior evidence of their sensitivity to environmental stressors such as humidity, hygiene, and confinement ^20^. While gut and oral niches remained resilient, these transient shifts at low-biomass skin sites may serve as early indicators of microbial perturbations, warranting targeted monitoring during longer missions. Additionally, we provide novel evidence of ISS-associated microbial post-flight emergence, interpersonal microbial transmission, and subject-specific bacterial exclusivity, underscoring the complexity of microbiome dynamics in confined habitats. In addition, the confined and shared environment of the ISS, including close interpersonal interactions among crew members, may facilitate microbial exchange and contribute to the observed variability in microbiome composition across individuals and body sites.

Gut microbiomes demonstrated strong resilience under spaceflight conditions, dominated by core taxa such as *Bacteroides* and *Prevotella*, consistent with terrestrial studies highlighting the stability of these communities in healthy adults ^20–22^. Subject-specific relative abundance patterns, including those observed for *Akkermansia* in Ax-2 and mutually exclusive *Prevotella* in Ax-3, suggest individualized metabolic adaptations rather than uniform shifts ^22^. Terrestrial studies have shown that gut microbiota maintains functional robustness despite environmental stressors ^14,23,24^. Prior work, including the NASA Twins Study, reported compositional changes during year-long missions that largely returned to baseline within six months post-flight ^15^ reinforcing the robustness of core gut communities. Our findings extend this evidence to short-duration commercial missions.

Urinary microbiomes exhibited low diversity and minimal temporal changes. In Ax-2, two participants displayed profiles dominated by *Lactobacillus* (up to 98%), while others showed higher relative abundances for *Streptococcus* and *Staphylococcus* (up to 82% and 87%). In contrast, Ax-3 urine samples were characterized by high proportions of *Sphingomonas* (up to 89%) and intermittent surges of *Staphylococcus* (up to 92%). In Ax-2, *Lactobacillus* dominance in two of the Ax-2 subjects is a noteworthy finding because there were two female subjects in that mission and because *Lactobacillus* has been reported to be associated with urine microbiome in females ^25^.

Skin-associated microbiomes showed the greatest sensitivity to spaceflight conditions, consistent with evidence from prior terrestrial studies indicating that these communities are highly dynamic and responsive to environmental stressors ^10,26,27^. Terrestrial studies have also shown that skin microbiomes respond to environmental stressors like humidity and hygiene changes; our findings reveal unique patterns under microgravity, including the emergence of opportunistic genera such as *Lawsonella* and *Sphingomonas*, which are rarely observed in Earth-based conditions. Across both missions, body swabs were dominated by *Corynebacterium* and *Staphylococcus*, taxa recognized as core commensals in maintaining skin barrier integrity ^26^. However, Ax-3 additionally exhibited opportunistic genera such as *Lawsonella* and *Sphingomonas*, which are not typically abundant in terrestrial skin microbiomes ^10^, revealing unique patterns under microgravity.

Taken together, these findings indicate that short-duration spaceflight exerts minimal impact on core microbial communities in saliva and stool ^14,28^, while urinary microbiomes remain stable with sex-dependent patterns ^25^. In contrast, skin-associated microbiomes display pronounced variability and opportunistic taxa, underscoring the need for targeted monitoring of low-biomass sites as early indicators of spaceflight-induced microbial shifts ^10,26^

Overall, these taxonomic patterns highlight strong subject-specific and body-site-specific microbial signatures across both missions. While certain taxa exhibited variability across time points and missions, these observations were not uniform across individuals and should be interpreted as descriptive trends rather than consistent spaceflight-driven effects. Given the limited cohort size, these findings are intended to provide a preliminary view of microbial dynamics and to guide future studies investigating taxonomic contributions to host responses. These patterns further underscore the importance of individualized microbiome responses as a key feature of the dataset and highlight the potential value of personalized monitoring approaches for understanding host-microbiome dynamics during spaceflight.

Temporal variation in the relative abundance of several microbial taxa was observed across missions and body sites, although these patterns were not uniformly consistent across individuals. To further assess these dynamics, we performed genus-level differential abundance analyses using MaAsLin3, accounting for repeated sampling within astronauts. These analyses identified a limited number of taxa with exploratory with exploratory associations with time point (q-value < 0.3) in selected body sites, providing additional support for a subset of the trends observed in descriptive taxonomic profiles. Complementary non-parametric Friedman tests similarly detected subtle within-subject temporal changes across sample types. Together, these results suggest that while temporal microbial shifts occur during and after spaceflight, they are generally subtle and may be influenced by strong inter-individual variability. Given the limited sample size, these associations should be considered hypothesis-generating observations.

Post-flight samples revealed rare, subject-specific bacterial emergence of low-abundance taxa, such as *Oligella* from subject SO4Fh9Mk9N (ARM) body swab samples (Ax-3 mission), which was further supported by WGS, providing high-resolution confirmation of the observation. Results suggest potential post-flight microbial emergence. WGS-based strain-level analysis identified the presence of two species, *Oligella urethralis* and *Oligella ureolytica*, in post-flight samples from this individual, with negligible or absent representation in pre-flight datasets. The marked increase in read counts at R+1, combined with near absence at pre-flight time points, support a post-flight emergence event.

To further support species-level classification, reads were evaluated using Kraken2, a k-mer–based taxonomic classifier. This analysis confirmed the presence of both *O. urethralis* and *O. ureolytica*-associated reads within the sample, providing complementary evidence for the detection of multiple *Oligella* species. These Gram-negative, non-fermentative coccobacilli are primarily associated with the human urogenital tract and are generally considered low-virulence organisms, although opportunistic infections have been reported ^29–31^. Because *Oligella* species are not commonly reported as environmental contaminants and are predominantly host-associated, their detection in a post-flight skin sample represents an unusual finding. This may reflect transient colonization, redistribution from host-associated reservoirs, or exposure within a shared confined environment such as the ISS. These findings, supported by WGS, suggest that microbial introductions were limited, individualized, and low in abundance rather than widespread across subjects.

Our study identified subject-specific bacterial taxa across sample types, showing potential personalized microbial signatures during spaceflight. Unique taxa included *Anaeroglobus* in saliva, *Anaeroplasma*, *Desulfovibrio*, and *Holdemanella* in stool, and *Holdemanella* across multiple skin sites, reinforcing individuality in microbiome profiles.

Post-flight elevations of IL-32 and IL-16 transcript expression in Ax-2 and Ax-3 suggest transient immune activation. Correlation analyses further indicated that these cytokines were associated with shifts in multiple microbial taxa across sample types, suggesting their potential involvement in observed host-microbiome interaction patterns. However, these associations are correlative, and additional studies are needed to determine whether direct mechanistic relationships exist between cytokine changes and microbial community shifts. IL-32 is a multifunctional pro-inflammatory cytokine that promotes innate and adaptive immune responses, contributes to epithelial barrier regulation, and has been implicated in host defense signaling under conditions of physiological stress ^9^. IL-16 functions as an immune chemoattractant that regulates T-cell activation, recruitment, and inflammatory signaling, serving as a mediator of immune surveillance and tissue homeostasis ^8^. These cytokines regulate inflammation, barrier integrity, and T-cell responses ^8,9,32^. The observation that the transcript expression of both cytokines increased following spaceflight in two independent missions supports the biological validity of the immune signatures identified in this study, as their established roles are consistent with the immune perturbations expected in response to microgravity, confinement, altered microbial exposures, and re-entry associated physiological stress ^1–3^. Although causality cannot be inferred from the present dataset, the reproducible post-flight increases observed across both missions strengthen the interpretation that the measured molecular responses reflect biologically meaningful physiological adaptation rather than technical variation. Longitudinal monitoring of cytokines such as IL-32 and IL-16 could enhance astronaut health surveillance by identifying individuals experiencing heightened immune activation or delayed recovery after flight. Inflight sampling remains logistically challenging and often incomplete, leaving gaps in understanding how and when microbiome changes occur. To address this, we propose implementing inflight collections using dry body swabs taken just before returning, with the swab collection buffer added immediately after splashdown to preserve samples. This approach mitigates the challenges of opening liquid-containing tubes in microgravity and enables safer, more practical sampling during future missions.

In conclusion, this study provides a comprehensive characterization of microbiome dynamics and immune responses during short-duration commercial spaceflight missions, revealing novel insights into microbial resilience, site-specific patterns. By integrating microbiome sequencing with cytokine transcript profiling, this study demonstrates that spaceflight induces coordinated shifts in skin-associated microbiomes and immune signaling, while core gut and oral communities remain largely stable. This study identified an instance of post-flight microbial emergence of environmental taxa from the spaceflight/ISS underscoring the complexity of host-microbe interactions in space. This work establishes a standardized framework for microbiome monitoring and immune surveillance applicable across multiple commercial spaceflights, to ensure astronaut safety on missions to the Moon, Mars, and beyond.

Collectively, this study provides several novel contributions to the field of space biology. First, the inclusion of multiple anatomical sampling sites across saliva, stool, urine, and skin-associated locations highlights strong subject-specific and site-specific microbial signatures, including instances of apparent microbial exclusivity. Second, the detection of low-abundance taxa in post-flight samples suggests the possibility of environmental or ISS-associated microbial emergence, underscoring the complexity of microbial exchange in confined habitats. Third, the integration of microbiome profiling with single-cell immune data enables a systems-level perspective linking microbial dynamics with elevated cytokines such as IL-32 and IL-16, at the transcript level. Finally, this work establishes a practical framework for longitudinal microbiome monitoring across missions and sample types, which can be extended to future commercial and long-duration spaceflight studies.

### Limitations of the study

The cohort size was inherently limited to six astronauts across two independent missions. Although longitudinal sampling generated a substantial number of biospecimens across multiple body sites and time points, the small number of participants limits statistical power and generalizability to broader astronaut populations.

Differences between missions should be interpreted cautiously. Unequal cohort distribution, along with variation in mission duration, environmental exposures, diet, physiology, sex-specific factors, operational conditions, and undocumented hygiene practices, limits the ability to distinguish mission-specific effects from inter-individual variability. Accordingly, observed differences in microbial abundance are considered exploratory and hypothesis-generating rather than definitive evidence of duration-associated effects.

Additionally, 16S rRNA gene amplicon sequencing provides genus-level resolution and variation in rRNA gene copy number may influence relative abundance estimates. Future studies integrating shotgun metagenomics, metatranscriptomics, and larger multi-omics cohorts will help to better define microbial functional adaptations and host interactions during spaceflight.

Although cytokine profiling at the transcript level provided complementary host-response context, the observational design precludes causal inference of microbiome-immune interactions. These analyses were not designed or powered to establish mechanistic relationships, so the associations between microbial and immune changes should be interpreted cautiously.

Interpretation of observed taxonomic shifts was guided by known ecological and functional roles, but these relationships warrant further validation in larger, controlled studies.

## Resource availability

### Lead Contact

Requests for further information and resources should be directed to and will be fulfilled by the lead contact, Harsha Doddapaneni

### Materials availability

This study did not generate any new unique reagents or materials.

### Data and code availability

- **Data:** Sequenced datasets from Microbiome and Single-cell sequences have been uploaded to the TRISH EXPAND TrialX database^33^ under the respective Microbiome and Single-cell data directories for the Axiom 2 and Axiom 3 missions. The data requests can be initiated through the TRISH EXPAND database portal ^33^. The TRISH EXPAND team enables data dissemination to the authorized researchers through an approved request process. Access to that data is governed by the EXPAND Database Privacy and Release Board (DPRB). The source data used to generate the figures presented in this manuscript can be accessed from GitHub (https://github.com/qiaoyanw/Olink-Explore-HT-Analysis-Methods-for-TRISH_AX2_AX3-Flight/tree/main)^34^.
- **Code:** No code was generated in this study.
- **All other items:** For any additional information, please contact.

## Acknowledgements

We thank Nathanael Emerick for his valuable assistance with the correlation analysis. This study was funded (Grant# INN0010) by the Translational Research Institute for Space Health through NASA Cooperative Agreement NNX16AO69A. Single-Cell libraries were prepared by the Single-Cell Genomics Core at Baylor College of Medicine, which is supported with funding from the CPRIT RP200504 and P30CA125123. The authors are grateful to the study participants and production teams at the Human Genome Sequencing Center for data generation.

## Author contributions

Conceptualization: H.D, R.A.G, A.K.

Data curation: T.M.S.R, M.C.R, A.K, Q.X, S.V.B, K.K, A.S, M.M.M, Y.C, M.C.G, Z.M, D.B. Formal analysis: A.K, T.M.S.R, Q.X, A.S, K.W, Q.W, D.K.

Writing-original draft: H.D, A.K.

Writing-review & editing: A.K, T.M.S.R, Z.M, A.S, S.V.B, K.K, K.W, D.K, E.U, S.M.G, M.M.M, J.W, D.M, R.A.G and H.D.

## STAR★METHODS

- KEY RESOURCE TABLE
- EXPERIMENTAL MODEL AND STUDY PARTICIPANT DETAILS

- Human study participants
- IRB statement on human subjects research
- METHODS DETAILS

## EXPERIMENTAL MODEL AND STUDY PARTICIPANT DETAILS

### Human study participants

Participant demographics are reported at the cohort level in accordance with IRB protocol BCM H-52035. Ax-2 participants (n = 4) ranged from 31 to 67 years (2 males and 2 females). For Ax-3, biospecimens were available from 2 of 4 crew members (n=2); additional demographic details are not reported to protect participant confidentiality.

### IRB statement on human subjects research

Protocol title: EXPAND-MESH (Enhancing exploration platforms and analog definition - multimodal evaluation of spaceflight participant health. (H-52035). Human biospecimens used in this study were obtained under protocols approved by the Institutional Review Board at Baylor College of Medicine (Protocol No. H-52035). All participants provided written informed consent for the collection and research use of their specimens. Samples were de-identified prior to analysis, and all research activities were conducted in accordance with institutional guidelines and applicable regulations.

## METHOD DETAILS

### Biospecimen collection and processing

In this study, data from two missions, that included a total of six astronauts (n = 6), and multiple biospecimens, across four sample types including saliva, stool, urine and body swabs, were collected according to the GENESTAR manual ^35^. A combination of low (body swabs, saliva, and urine) and high biomass (stool) samples (Figure 1A) was collected from 4 subjects from Ax-2 mission, and 2 subjects from Ax-3 mission. Body swab samples were collected from 10 different body locations including post-auricular (EAR), axillary vault (PIT), volar forearm (ARM), occiput (NAP), umbilicus (UMB), gluteal crease (GLU), glabella (TZO), toe web space (WEB), oral swab (ORC), and nasal swab (NAC) (Figure 1A).

Ax-2 samples were collected at 30-and 3-days pre-launch (L-30, L-3), and 6 days after returning (R+6) from the ISS. Samples collected at L-90 from Ax-2 were not utilized in this study. Ax-3 samples were collected at 30- and 2-days pre-launch (L-30, L-2), and 1 and 21 days after returning (R+1 and R+21) from the ISS (Figure 1B). A total of 11 saliva, 12 stool, 12 urine and 120 body swab samples collected from four participants from Ax-2; and 8 saliva, 8 stool, 8 urine samples, and 80 body swab samples collected from two subjects from Ax-3, which were then processed through 16S rRNA gene (V4 region) amplicon sequencing. One subject could not provide a saliva sample after their return from the ISS (Table S1).

### DNA extraction and 16S rRNA gene V4 region amplification

Stool, saliva, urine, and body swab samples totaling 155 from Ax-2 and 104 from Ax-3 were extracted using Qiagen DNeasy PowerSoil Pro Kit following manufacturer’s instructions. One sample from Ax-2 failed at the QC step and was excluded, leaving 154 samples from Ax-2 and 104 from Ax-3. Together, 258 samples proceeded for sequencing. The 16S rRNA gene V4 region was then amplified using the primers 515F (GTGCCAGCMGCCGCGGTAA) and 806R (GGACTACHVGGGTWTCTAAT). Primers used for amplification contain adapters for MiSeq sequencing and single-index barcodes incorporated into the reverse primer so that resulting PCR products that can be pooled and sequenced directly, targeting an average per sample yield of approximately 25,000 read pairs. Products were then sequenced on the Illumina MiSeq platform (2 x 250 bp).

### Data annotation

Raw data files in binary base call (BCL) format were converted into FASTQs and demultiplexed based on the single-index barcodes using the Illumina ‘bcl2fastq’ software. Demultiplexed read pairs underwent an initial quality filtering using bbduk.sh (BBMap, version 38.82), removing Illumina adapters, PhiX reads and reads with a Phred quality score <15 and length <100 bp after trimming. Quality controlled reads were merged using bbmerge.sh with merge parameters: maxstrict=t, qtrim=t, trimq=15. Merged reads were further filtered via vsearch using parameters optimized for the V4 region of the 16S rRNA gene. All reads were then combined into a single FASTA file for further processing using UPARSE. Resulting reads are clustered into OTUs at similarity cutoff 97% via the UPARSE algorithm using an in-house stepwise approach that includes chimera filtering. Briefly, reads are first dereplicated using the vsearch ‘derep_fulllength’ option, recording the size to identify singletons. Dereplicated non-singleton reads are clustered through an iterative stepwise manner in increments of 0.4% using the usearch70 ‘cluster_otus’ function. Singletons are mapped back to the dereplicated reads using usearch70 ‘-usearch_global’9 at a 99% identity and mapped reads are added to an output file. The final output is run through usearch70 ‘uchime_ref’ against the GOLD database v510.11, using only the positive-sense strand and allowing for no chimeras. The OTU centroids are then mapped against an optimized version of the latest current SILVA Database containing only sequences from the V4 region of the 16S rRNA gene to determine taxonomies using the usearch70 ‘usearch_global’ function9, specifying the identity threshold to 97%. Abundances are recovered by mapping demultiplexed reads to the OTU file, creating an OTU table in biom format and removing chimeric reads. Phylogeny information contained in the biom file was generated by aligning the centroid sequences with mafft and creating a tree via FastTree.

### Data analysis

Prior to analysis, rarefaction was performed on a per-sample type basis to minimize the effect of differing rarefaction cutoff values on diversity and taxonomic profiles. Once samples were rarefied, diversity and taxonomic analyses were performed, as described below. Taxonomic abundance visualizations were generated from relative abundance profiles within each sample type and mission, emphasizing the most abundant and consistently detected genera across longitudinal samples to characterize dominant microbial patterns. Given the cohort size, no formal prevalence filtering, differential abundance testing, or predefined selection criteria were applied. Relative abundance estimates were derived from normalized 16S rRNA gene amplicon sequencing data using standard workflows.

Because stool, saliva, urine, and skin-associated biospecimens differed substantially in microbial biomass and sequencing yield, sample type-specific rarefaction thresholds were applied to maximize longitudinal sample retention while minimizing exclusion of lower-biomass samples. Consequently, diversity analyses were primarily interpreted within biological sample categories, and cross-sample type or cross-mission quantitative comparisons should be approached cautiously. In addition, standardized biospecimen collection, processing, storage, and transport protocols were applied consistently across all participants and missions, minimizing technical variability and supporting the reproducibility of observed microbiome patterns, particularly for skin-associated samples.

### Diversity analysis

Diversity analyses were performed using ATIMA (https://atima.research.bcm.edu/), a stand-alone tool for analyzing and visualizing microbiome datasets. The software is a web application combining publicly available R packages with purpose-written code to import sample data and identify trends in taxa abundance, alpha diversity, and as they relate to sample metadata. Significant differences between variables in the alpha diversity analysis were evaluated using the non-parametric Mann-Whitney U test or Kruskal-Wallis for the comparison of 2 groups or 3 or more groups, respectively ^36,37^. Alpha diversity results were visualized through box plots, and p-values were adjusted using False Discovery Rate (FDR) correction ^38^. Beta diversity analyses were performed using the R function vegan:adonis version 2.5.5 to estimate Bray-Curtis distances and visualized through Principal Coordinate Analyses (PCoA) plots. Beta diversity significance was assessed through a Permutational Multivariate Analysis of Variance (PERMANOVA) ^39^.For within-subject longitudinal comparisons across time points, non-parametric Friedman tests were performed, followed by paired post hoc comparisons where appropriate. These approaches were selected to account for repeated measurements within individuals and the small cohort size.

### Taxonomic abundance and differential abundance

Multidimensional bubble plots were generated in RStudio (Posit, v2024.x) using R (v4.3.2) and the ggplot2 package to visualize bacterial genera across urine, saliva, and stool samples from Ax-2 and Ax-3 missions. Data were formatted in a long structure with columns for sample type, subject ID, mission, time point, genus, and relative abundance (%). The top 10 most abundant genera per sample type were selected based on mean abundance across all time points. Bubble size was scaled to represent relative abundance (0–100%), and colors of distinguished sample types. Facet wrapping was applied to separate subjects and missions, enabling simultaneous comparison of temporal trends and subject-specific patterns. Additional customization included axis ordering for time points, legend annotations for abundance scaling, and theme adjustments for clarity. This visualization provided an integrated view of microbial composition dynamics across multiple biological niches.

Temporal associations between microbial taxa and mission time point were evaluated using MaAsLin3 ^40^. Genus-level relative abundance tables were analyzed separately for each body site. Low-prevalence taxa were filtered prior to analysis, and data were normalized using total sum scaling (TSS) and log transformation. Microbial abundance was modeled as a function of time point while accounting for repeated sampling using subject-specific random effects. For both missions, L-30 served as the reference time point (Ax-2: L-3, R+6; Ax-3: L-2, R+1, R+21). P-values were adjusted using the Benjamini-Hochberg method, with q-values < 0.30 considered significant for exploratory analyses and hypothesis-generating.

### Whole Genome Sequencing

Whole genome sequencing was performed, utilizing the DNA prepared from Ax-2 and Ax-3 missions, on the 4 samples (Table S2) to verify the results from 16S rRNA gene amplicon sequencing. Illumina paired-end libraries were prepared and sequenced on NovaseqX 25B flow cell (2×150bp reads) (Table S2).

Quality and adaptor trimming for Illumina WGS reads was performed using BBMap version 39.01 (https://github.com/BioInfoTools/BBMap). After removing human host reads using BBMap, taxonomic profiling of WGS samples was conducted using MetaPhlAn version 4.2.4 (https://github.com/biobakery/MetaPhlAn). Kraken2 version 2.1.6 (https://github.com/DerrickWood/kraken2) was also used for taxonomic classification for WGS reads in comparison to the results from MetaPhlAn and 16S rRNA sequencing.

### PBMC Isolation and Single-cell RNA workflow

Blood samples were collected at L-30, L-3, and R+1 for Ax-2 (n = 4) and L-30, L-2, and R+1 for Ax-3 (n = 2), yielding 17 samples for single-cell RNA sequencing. PBMCs were isolated using standard GENESTAR protocols, with viability assessed by trypan blue prior to cryopreservation. Single-cell 5′ gene expression libraries were generated using the 10x Genomics Chromium platform (v2 for Ax-2; v3 for Ax-3) and sequenced on a NovaSeq X system. Data were processed using Cell Ranger (GRCh38 reference). Single-cell Analysis Data were analyzed using Seurat (v5) following standard workflows for quality control, normalization, dimensionality reduction, clustering, and integration. Cell populations were annotated based on marker genes, and cytokine expression patterns were assessed using differential expression analysis. Detailed workflows are described in the companion manuscript.

### Cytokine Measurement from Blood Samples

Blood samples were collected at time points L-30, L-3, and R+1 from four subjects, and L-30, L-2, and R+1 from two subjects during the Ax-2 and Ax-3 missions. A total of 17 samples were used for single-cell RNA sequencing to assess cytokine expression. Details of these datasets can be found in the companion paper ^19^.

### Correlation analysis

To assess associations between microbiome dynamics and host immune responses, we performed correlation analyses between changes in microbial taxonomic abundance and circulating cytokine transcript levels in the Axiom 2 mission, the only dataset yielding analyzable results (Table S3). Microbial relative abundances were CLR-transformed to account for compositionality, and changes were defined per subject as post-flight (R+6) minus pre-flight baseline (L-30). Cytokine responses (IL-16 and IL-32) were calculated as early post-flight (R+1) minus baseline (L−30). Because measurements were collected at slightly different post-flight time points, comparisons reflect overall post-flight changes rather than exact matched-timepoint differences. Pearson correlation was used to assess linear relationships, with nominal significance defined as p values < 0.05 are reported (Table S3) along with subject-level correlation analysis shown in Figure S7-S9. Results may be sensitive to individual subjects. Positive correlations should be interpreted as associations between relative shifts in microbial composition and cytokine transcript responses rather than absolute changes, but useful for hypothesis generation. Correlation analysis was not performed for Axiom 3 due to insufficient sample size (n = 2).

### Quantification and statistical analysis

Alpha diversity was evaluated using Mann-Whitney U or Kruskal-Wallis tests, as appropriate, with FDR correction for multiple comparisons. Beta diversity was assessed using PERMANOVA based on Bray-Curtis distances. Longitudinal within-subject comparisons were evaluated using Friedman tests followed by paired post hoc comparisons where appropriate. Differential abundance was assessed using MaAsLin3 with Benjamini-Hochberg correction, with q < 0.30 considered significant for exploratory analyses. Pearson correlation was used for microbiome-interleukin associations, with nominal significance defined as p < 0.05. There was no reportable quantification analysis performed in this study.

### Supplemental information

**Document S1. Figures S1-S9, Tables S1-S3.**

