## Supplemental Table 3 for "Microbial Dynamics Across Commercial Spaceflights of Varying Duration"

**Figure S1**

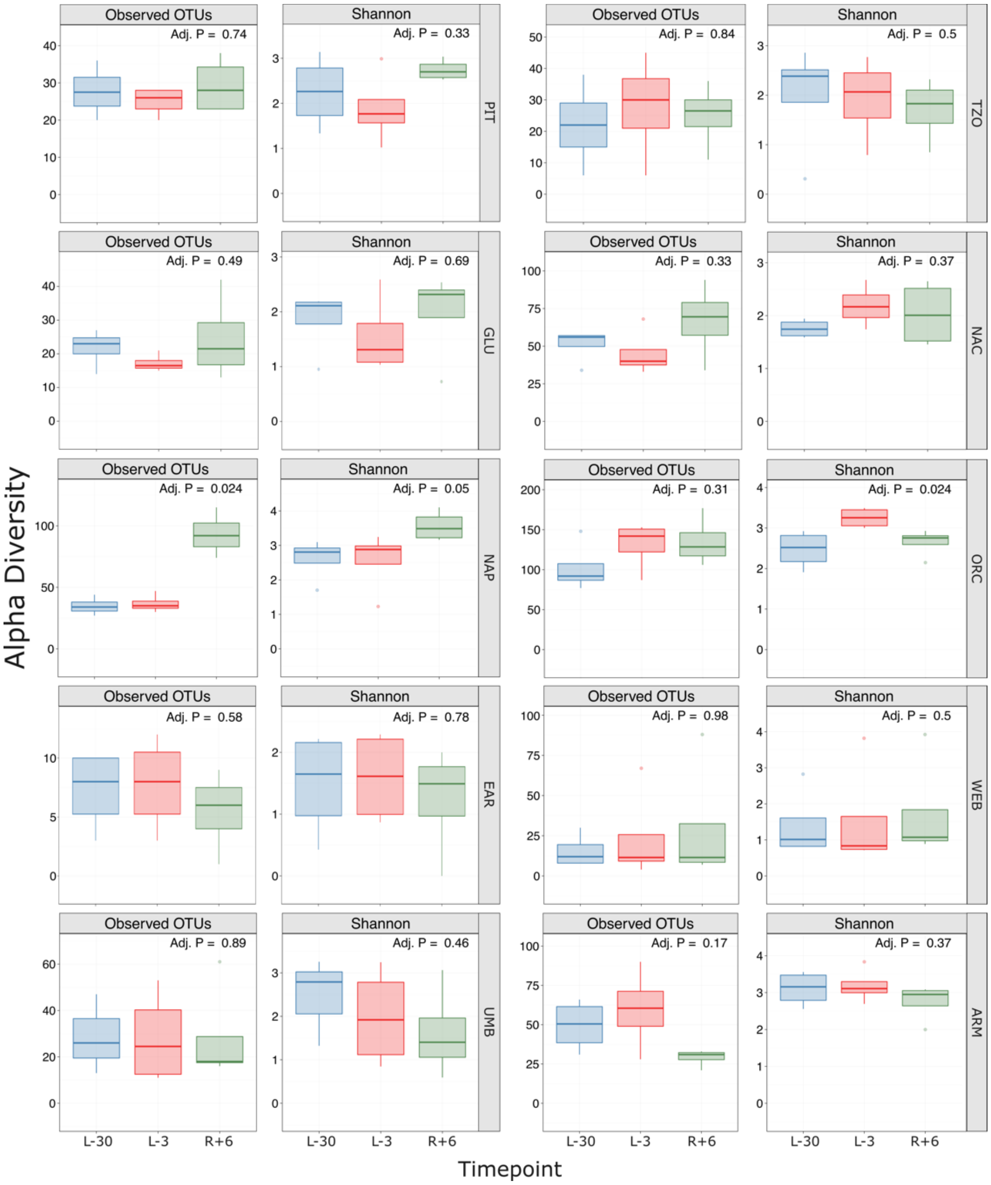


**Figure S1. Alpha diversity analysis of body swab samples from Ax-2.** The figure shows the number of observed Operational Taxonomic Units (OTUs) and Shannon diversity index for body swab samples, including PIT, TZO, GLU, NAC, NAP, ORC, EAR, WEB, UMB, and ARM. Boxplots are colored by timepoints, namely L-30, L-3, and R+6. Adjusted p-values are shown on the plots.

**Figure S2**

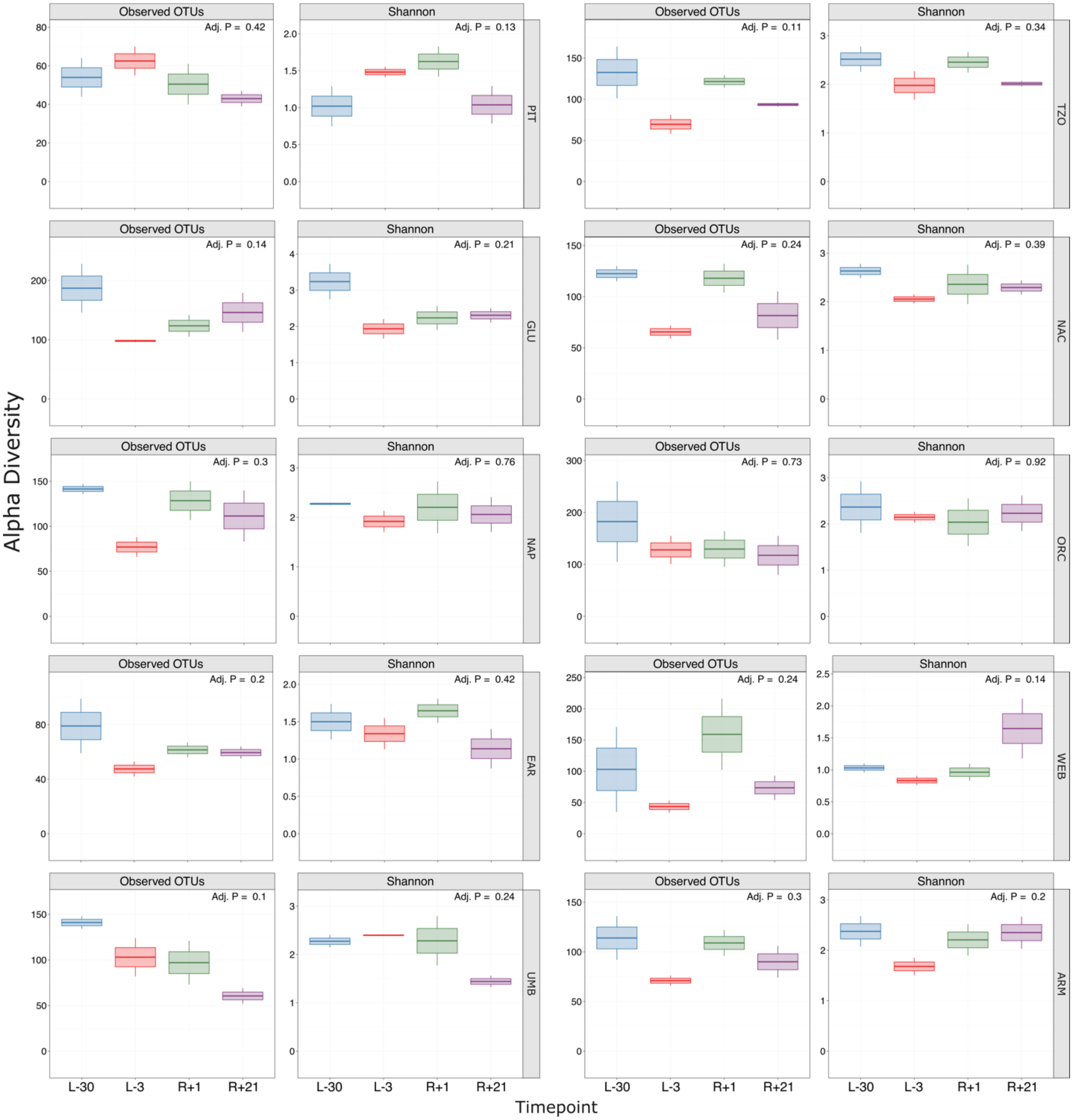


**Figure S2. Alpha diversity analysis of body swab samples from Ax-3.** The figure shows the number of observed Operational Taxonomic Units (OTUs) and Shannon diversity index for body swab samples, including PIT, TZO, GLU, NAC, NAP, ORC, EAR, WEB, UMB, and ARM. Boxplots are colored by timepoints, namely L-30, L-3, R+1 and R+21. Adjusted p-values are shown on the plots.

### Figure S3

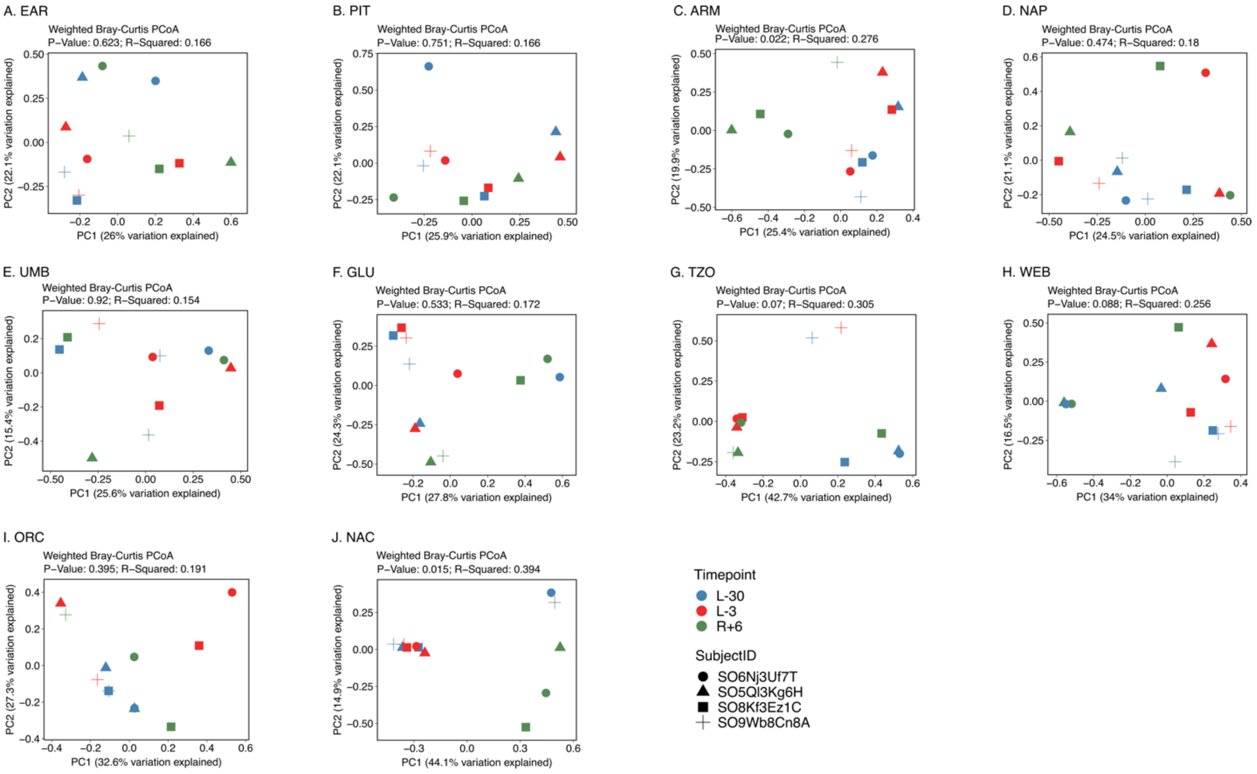


**Figure S3. Beta diversity analysis of body swab samples collected from the Ax-2 mission.** Results show PCoA plots of the Bray-Curtis distances of EAR, PIT, ARM, NAP, UMB, GLU, TZO, WEB, ORC, and NAC samples (Panels A-J) samples across subjects and timepoints. P-values were calculated using PERMANOVA and are displayed on the plot along with the R-squared value.

### Figure S4

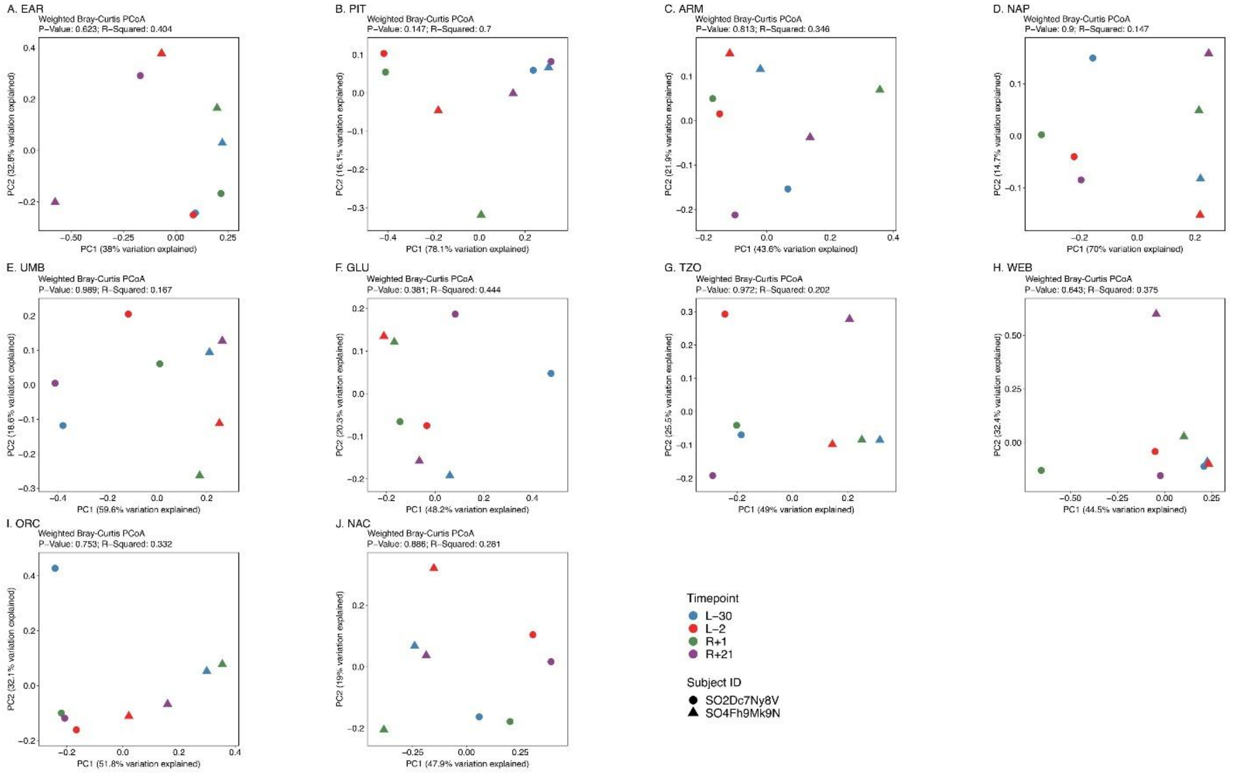


**Figure S4. Beta diversity analysis of body swab samples collected from the Ax-3 mission.** Results show PcoA plots of the Bray-Curtis distances of EAR, PIT, ARM, NAP, UMB, GLU, TZO, WEB, ORC, and NAC (Panels A-J) samples across subjects and timepoints. P-values were calculated using PERMANOVA and are displayed on the plot along with the method, R-squared, and F-statistic values.

### Figure S5

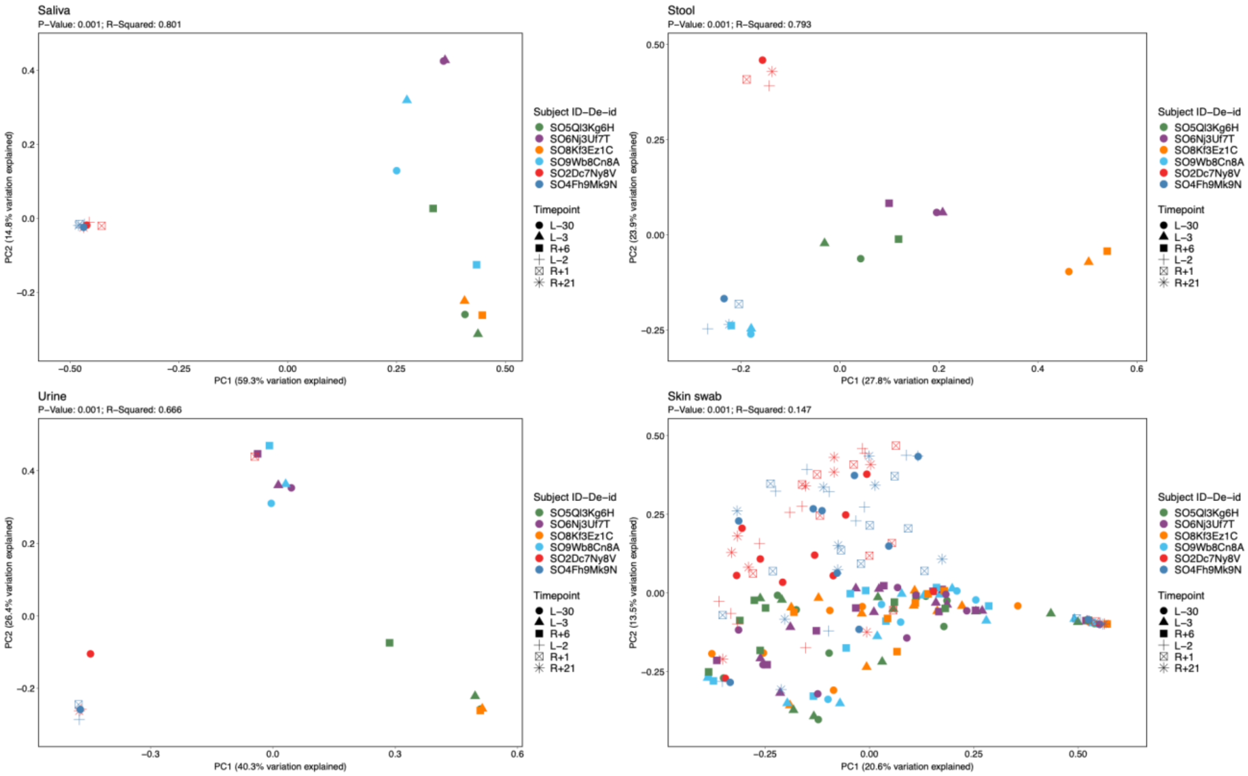


**Figure S5:** Weighted Bray-Curtis Principal Coordinate Analysis (PCoA) plots faceted by biological sample type across the Axiom-2 and Axiom-3 missions. Saliva, stool, urine and body swab microbiomes are displayed separately to evaluate longitudinal microbial dynamics independent of dominant body site-associated clustering. Points are colored by subject ID and shaped by timepoint. PERMANOVA P-values and $R^2$ values are shown within each panel.

**Figure S6**

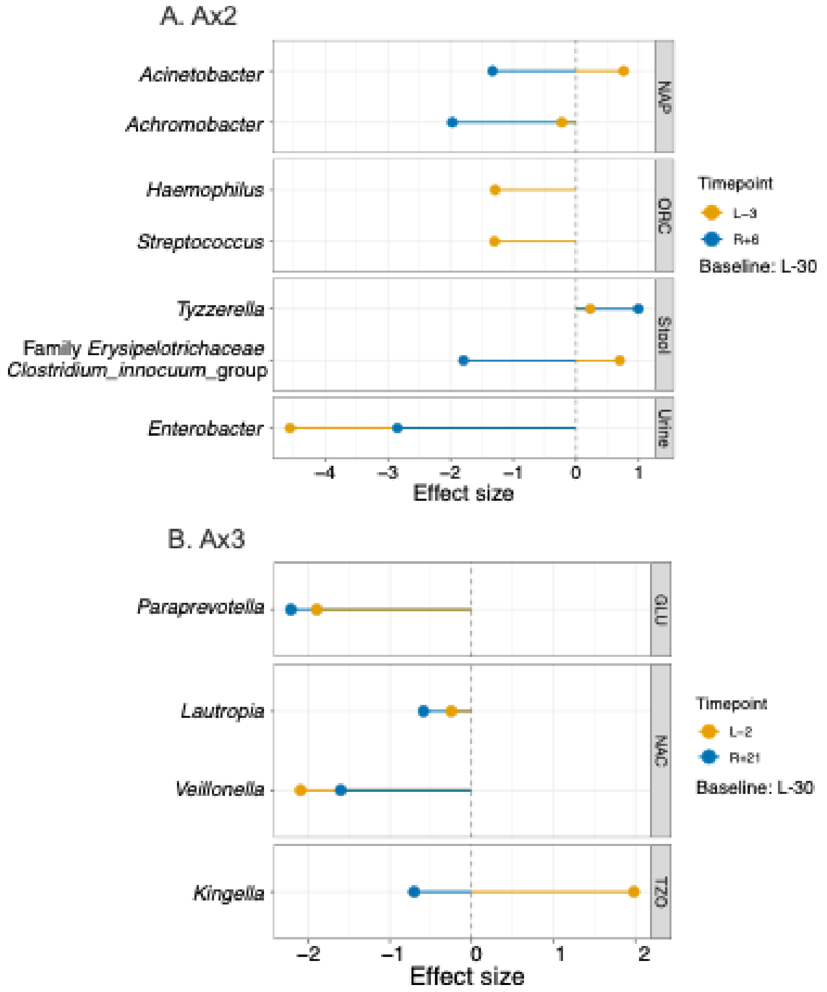


**Figure S6:** Longitudinal changes in microbial taxa across the Ax-2 and Ax-3 missions. (A) Ax-2 and (B) Ax-3 genera significantly associated with sampling timepoint identified using MaAsLin3 mixed-effects models. For each body site, microbial abundance was modeled as a function of timepoint while accounting for repeated measurements within astronauts using subject-specific random effects. Lollipop plots display model coefficients (effect sizes) for taxa meeting the significance threshold ($q < 0.25$). Positive coefficients indicate higher relative abundance compared with the pre-flight baseline timepoint, whereas negative coefficients indicate lower relative abundance. Colors denote the timepoint comparison relative to baseline (Ax-2: L-3 and R+6; Ax-3: L-2 and R+21), and point shapes indicate whether the association was detected in abundance- or prevalence-based models. Facets represent body sites with significant associations.

### Figure S7

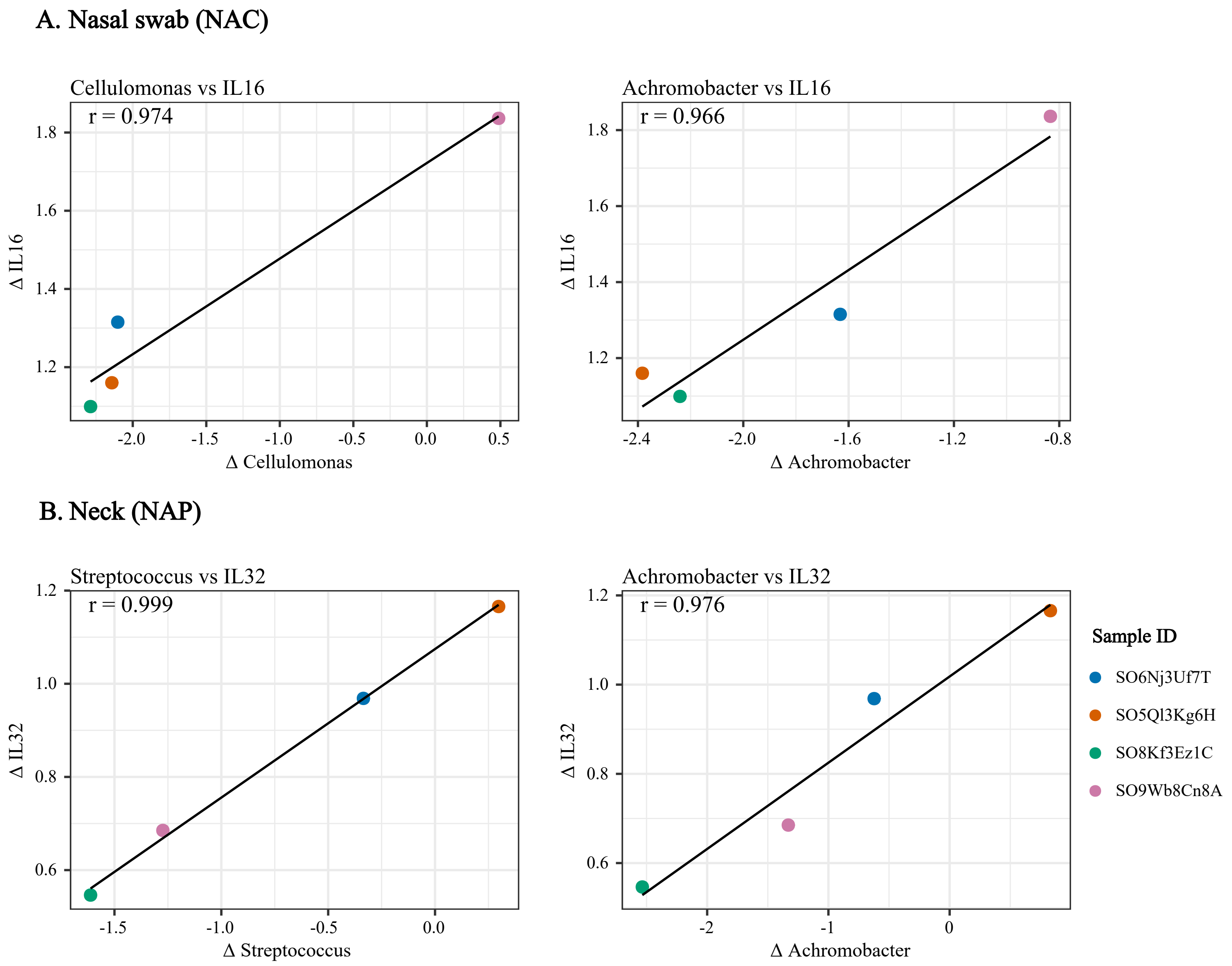


**Figure S7.** Subject-level microbiome-cytokine correlation analyses in the Ax-2 cohort (n = 4). Scatterplots showing correlations between subject-specific post-flight versus baseline changes in microbial relative abundance and cytokine (IL-16, IL32) transcript expression, from nasal swab (NAC), and neck (NAP) samples, summarized in Table S3. Each point represents an individual participant. Pearson correlation coefficients (r) are shown for all 4 associations. (A) Nasal swab and (B) Neck.

**Figure S8**

**Oral swab (ORC)**

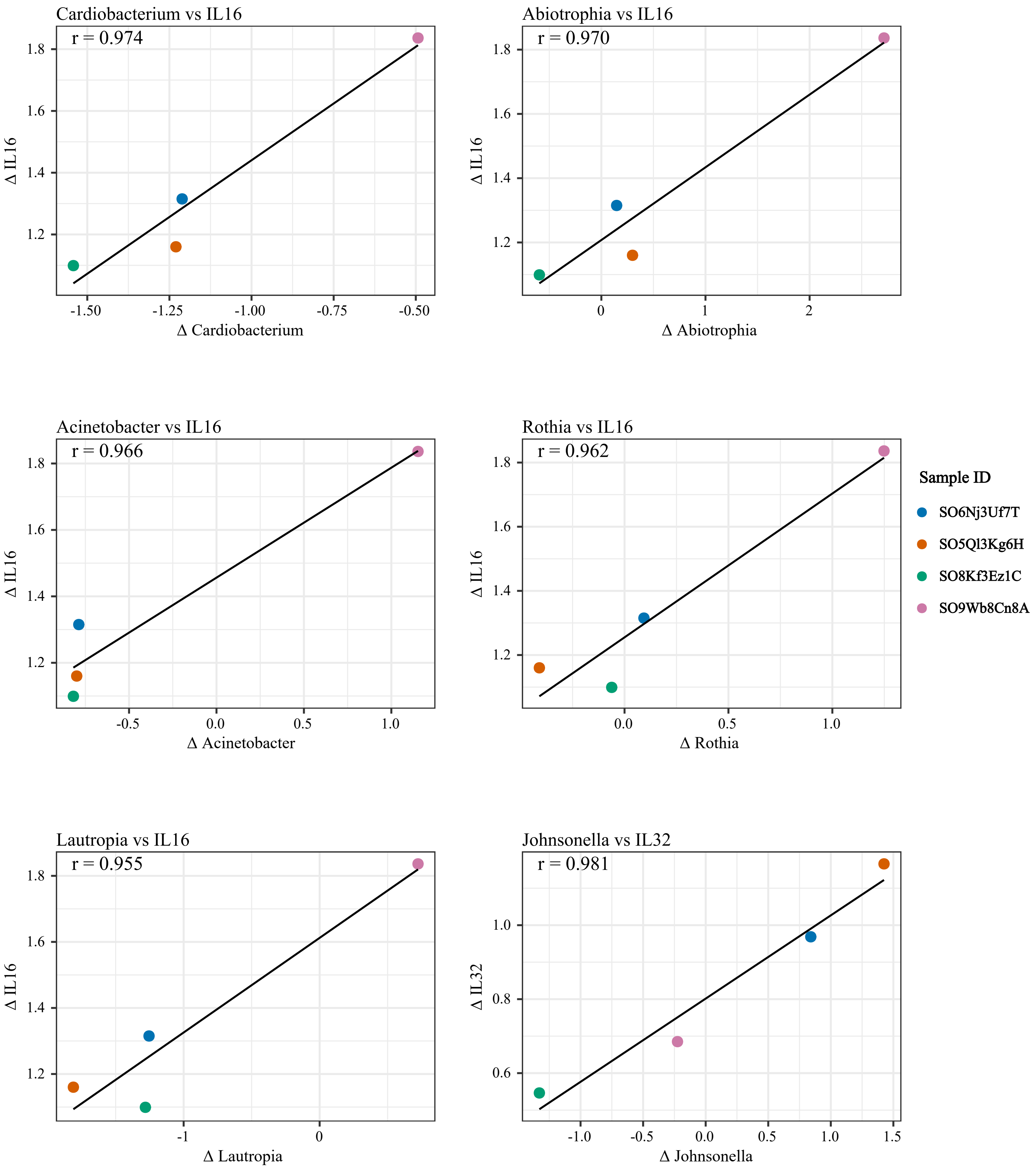


**Figure S8.** Subject-level microbiome-cytokine correlation analyses in the Ax-2 cohort (n = 4). Scatterplots showing correlations between subject-specific post-flight versus baseline changes in microbial relative abundance and cytokine (IL-16, IL32) transcript expression, from oral swab (ORC) samples, summarized in Table S3. Each point represents an individual participant. Pearson correlation coefficients (r) are shown for all 6 associations.

### Figure S9

#### A. Stool

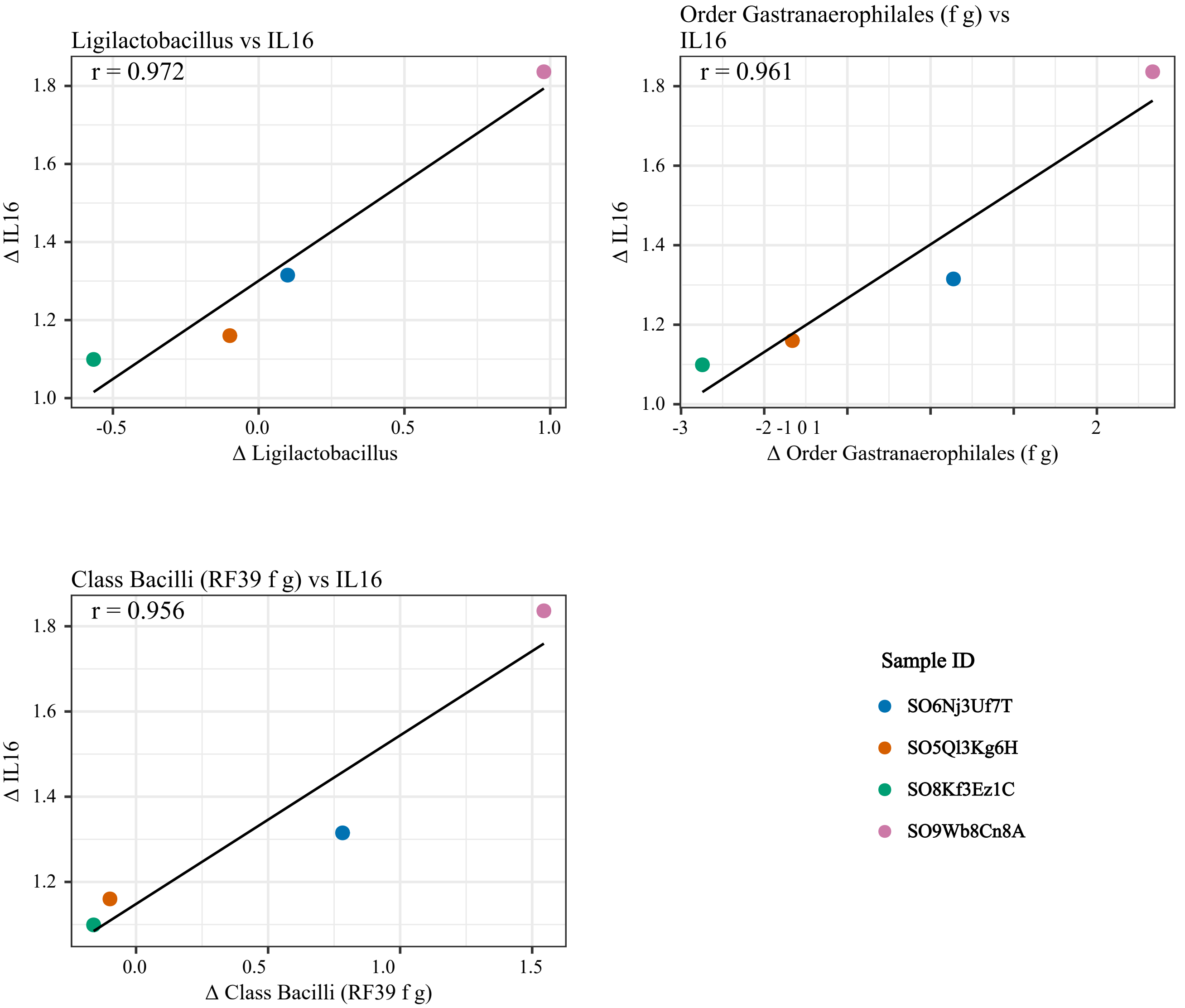


#### B. Urine

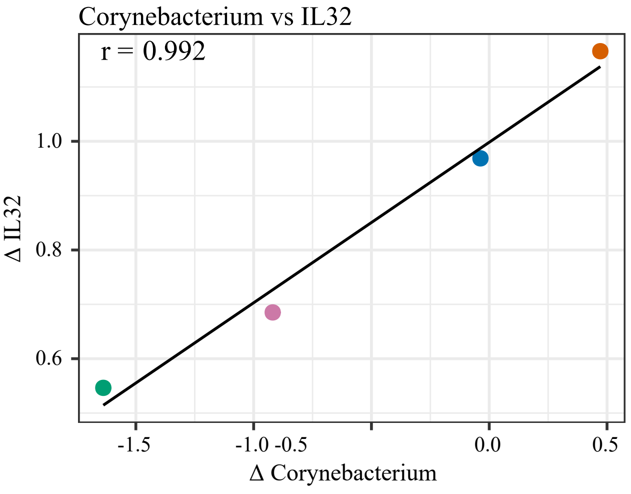


**Figure S9**. Subject-level microbiome-cytokine correlation analyses in the Ax-2 cohort (n = 4). Scatterplots showing correlations between subject-specific post-flight versus baseline changes in microbial relative abundance and cytokine (IL-16, IL32) transcript expression, from stool and urine samples, summarized in Table S3. Each point represents an individual participant. Pearson correlation coefficients (r) are shown for all 4 associations. (A) Stool and (B) Urine.

**Table S1.** Sample description (Related to Figure 1).

| | Sample type | Description | Body swab type | Swab collection description | Sample # from each mission | |
|---|---|---|---|---|---|---|
| | | | | | Axiom 2 | Axiom 3 |
| | Stool | | | | 12 | 8 |
| | Saliva | | | | 11 | 8 |
| | Urine | | | | 12 | 8 |
| Body Swab | Post-Auricular (EAR) | Behind the ear | Wet swabs | Swabs will be first dipped in sterile nuclease-free water before contact with the skin | 12 | 8 |
| | Axillary Vault (PIT) | Armpit | | | 12 | 8 |
| | Volar Forearm (ARM) | Forearm | | | 12 | 8 |
| | Occiput (NAP) | Upper back/Nape of neck | | | 12 | 8 |
| | Umbilicus (UMB) | Belly button | | | 12 | 8 |
| | Gluteal Crease (GLU) | Upper crack | | | 12 | 8 |
| | Glabella (TZO) | T-zone | | | 12 | 8 |
| | Toe Web Space (WEB) | Between the toes | | | 12 | 8 |
| | Oral Swab (ORC) | Side of cheeks | Dry swabs | Swabs will be dry upon skin contact | 12 | 8 |
| | Nasal Swab (NAC) | Lower nasal Cavity | | | 12 | 8 |

**Table S2. Bacterial verification**. This table presents bacterial read counts for *Akkermansia* and *Oligella* across pre- and postflight samples from subjects SO8Kf3Ez1C and SO4Fh9Mk9N, respectively. It includes data from 16S rRNA gene sequencing and whole genome-genome sequencing (WGS), total sequencing depth, bacterial read assignments, and associated sample metadata.

| Bacterial genera | Subject ID | Time point | Sample type-location | Mission | Category | Bacterial read counts | Sequencing depth | Bacterial read counts | Sequencing depth | Number of reads mapped to Bacteria (Kraken2) |
|---|---|---|---|---|---|---|---|---|---|---|
| | | | | | | 16S rRNA gene | | WGS | | |
| *Akkermansia* | SO8Kf3Ez1C | L-30 | Stool | Axiom 2 | | 0 | 72,727 | NA | | |
| *Akkermansia* | SO8Kf3Ez1C | L-3 | Stool | Axiom 2 | Significant post-flight surge | 2 | 76,653 | 402 | 29,533,596 | 15,177,350 |
| *Akkermansia* | SO8Kf3Ez1C | R+6 | Stool | Axiom 2 | | 10340 | 84,183 | 288,364 | 27,287,724 | 16,914,078 |
| *Oligella* | SO4Fh9Mk9N | L-30 | Body Swab - ARM | Axiom 3 | | 0 | 11,233 | NA | | |
| *Oligella* | SO4Fh9Mk9N | L-2 | Body Swab - ARM | Axiom 3 | Space-flight/ISS acquisition | 0 | 9,400 | 1 | 29,648,760 | 16,711,182 |
| *Oligella* | SO4Fh9Mk9N | R+1 | Body Swab - ARM | Axiom 3 | | 34 | 13,484 | 2904 | 34,218,186 | 17,142,212 |

Quality and adaptor trimming for Illumina WGS reads was performed using BBMap version 39.01 (https://github.com/BioInfoTools/BBMap). After removing human host reads using BBMap, taxonomic profiling of WGS samples was conducted using Kraken2 version 2.1.6 (https://github.com/DerrickWood/kraken2) to evaluate bacterial composition in comparison with results from 16S rRNA gene sequencing.

**Table S3. Correlations between microbial taxa and interleukin levels across sample types (Related to Figure S7).** This table summarizes associations between microbial taxa and cytokine concentrations (IL16 and IL32) across multiple sample types (nasal swab, neck, oral swab, stool, and urine). For each microbe–interleukin pair, correlations were calculated using both Pearson and Spearman methods across n = 4 subjects. “Method agreement” indicates concordance between Pearson and Spearman correlation directionality. Statistical significance is reported as raw p-values. Taxonomic annotations are reported at the most specific available level; entries labeled with “f g” denote family/genus-level classifications where full taxonomic resolution was not available.

| Sample Type | Microbe | Interleukin | n_subjects | Pearson_r | Spearman_r | method_agreement | p_value |
|---|---|---|---|---|---|---|---|
| Nasal Swab | Cellulomonas | IL16 | 4 | 0.974430251 | 1 | agree | 0.02556975 |
| Nasal Swab | Achromobacter | IL16 | 4 | 0.966141751 | 0.8 | agree | 0.03385825 |
| Neck | Streptococcus | IL32 | 4 | 0.998936281 | 1 | agree | 0.00106372 |
| Neck | Achromobacter | IL32 | 4 | 0.975520203 | 1 | agree | 0.0244798 |
| Oral Swab | Johnsonella | IL32 | 4 | 0.981311881 | 1 | agree | 0.01868812 |
| Oral Swab | Cardiobacterium | IL16 | 4 | 0.97428439 | 1 | agree | 0.02571561 |
| Oral Swab | Abiotrophia | IL16 | 4 | 0.970312822 | 0.8 | agree | 0.02968718 |
| Oral Swab | Acinetobacter | IL16 | 4 | 0.965680076 | 1 | agree | 0.03431992 |
| Oral Swab | Rothia | IL16 | 4 | 0.961966489 | 0.8 | agree | 0.03803351 |
| Oral Swab | Lautropia | IL16 | 4 | 0.954524922 | 0.8 | agree | 0.04547508 |
| Stool | Ligilactobacillus | IL16 | 4 | 0.972364422 | 1 | agree | 0.02763558 |
| Stool | Order Gastranaerophilales (f g) | IL16 | 4 | 0.961086958 | 1 | agree | 0.03891304 |
| Stool | Class Bacilli (RF39 f g) | IL16 | 4 | 0.955857602 | 1 | agree | 0.0441424 |
| Urine | Corynebacterium | IL32 | 4 | 0.991586895 | 1 | agree | 0.00841311 |

Data S1. Selected subject- and sample type–specific taxonomic patterns

**Saliva microbiome (subject-specific patterns)**

In subject SO6Nj3Uf7T, *Corynebacterium* declined progressively from 69.18% (L−30) and 49.51% (L−3) to undetectable levels by R+6. Subject SO8Kf3Ez1C exhibited a shift in dominance from *Streptococcus* (15.41% at L−30) to *Staphylococcus* at L−3 (52.13%) and R+6 (72.46%), with a concurrent increase in *Corynebacterium* to 19.34% at R+6. In contrast, SO9Wb8Cn8A displayed alternating dominance between *Staphylococcus* (26.23% at L−30 and 49.18% at R+6) and *Corynebacterium* (33.44% at L−3).

At the community level, oral microbiomes in Ax-3 were dominated by *Streptococcus* and *Veillonella* immediately post-flight and persisted through R+21, whereas Ax-2 samples approached baseline composition by R+6. Gut-associated trends indicated reductions in *Prevotella* and increases in *Faecalibacterium*, *Roseburia*, and *Blautia* through R+21 in Ax-3, compared to partial normalization by R+6 in Ax-2.

**Stool microbiome (sample-type patterns)**

Stool microbiomes remained dominated by gut-associated taxa while exhibiting mission-specific temporal variation. In Ax-2, subjects displayed distinct enrichment patterns, particularly for *Akkermansia*. Subject SO6Nj3Uf7T showed a high relative abundance at L−30 (41.76%), decreasing to 14.80% by R+6, whereas SO8Kf3Ez1C exhibited a pronounced post-flight (R+6) increase in *Akkermansia,* increasing from negligible to 17%. This transient increase was further supported by whole-genome sequencing (WGS) data (Table S2). Subject SO9Wb8Cn8A, in contrast, was enriched for *Prevotella_9*, with relative abundances of 27.44% at L−30 and 43.80% at R+6.

In Ax-3, stool microbiomes displayed subject-associated dominance patterns. Subject SO2Dc7Ny8V showed enrichment of *Prevotellaceae_NK3B31* (13.94% at L−2; 23.40% at R+21) alongside *Bacteroides* (14.62%–20.70%). In contrast, SO4Fh9Mk9N was dominated by *Prevotella_9* (18.28% at R+1; 37.21% at R+21) and exhibited a decline in *Faecalibacterium* (17.74% at L−30 to 7.09% at R+21).

**Urinary microbiome (subject- and mission-specific patterns)**

Urinary microbiomes demonstrated marked inter-individual and mission-specific variation. In Ax-2, two subjects exhibited strong dominance of *Lactobacillus*, comprising 90.22%–97.76% and 77.54%–97% across timepoints. In the remaining two subjects, urinary communities were more variable, with *Staphylococcus* (3.47%–87.27%) and *Corynebacterium* (3.12%–52.06%) representing the predominant taxa.

In contrast, Ax-3 urine samples were primarily dominated by *Sphingomonas*, with consistently high relative abundances across timepoints (63.14% at L−30, 84.22% at L−2, and 89.24% at R+21). Transient surges of *Staphylococcus* were observed (e.g., 91.71% at R+1), along with occasional detection of *Streptococcus*, indicating episodic fluctuations superimposed on a stable dominant background.

**Body swab microbiome (subject- and site-specific patterns)**

Body swab microbiomes reflected strong subject- and site-specific variation across both missions. In Ax-2, Corynebacterium (up to 95%) and Staphylococcus (up to 100%) were the dominant taxa, whereas Ax-3 additionally exhibited enrichment of *Lawsonella* (up to 50.89%) and *Sphingomonas* (up to 69.80%). Site-specific patterns were evident across subjects. In Ax-2, TZO samples were dominated by *Corynebacterium* in SO6Nj3Uf7T (75%–94.79%) and SO9Wb8Cn8A (46.88%–71.35%). In Ax-3, NAP samples showed marked enrichment of *Lawsonella* in SO2Dc7Ny8V (28.56% at L−30 increasing to 50.89% at R+21), compared to consistently low levels in SO4Fh9Mk9N (1.52%–3.81%). ORC samples consistently reflected oral-associated taxa, with *Streptococcus* (up to 51%) and *Haemophilus* (up to 33%) dominating post-flight timepoints. Opportunistic genera, including *Micrococcus* (up to 6%), *Paracoccus* (up to 2%), and *Comamonas* (up to 3%), were observed exclusively in post-flight samples.
